# Nucleophagy removes cytotoxic trapped PARP1

**DOI:** 10.64898/2025.12.11.692727

**Authors:** Gwendoline Hoslett, Sara Tribble, Pauline Lascaux, Stelios Koukouravas, Ignacio Torrecilla, Wei Song, Cynthia Hou, Junyi Li, Martín González-Fernández, Giuliana De Gregoriis, Rebecca A Dagg, Darragh O’Brien, Andrea Pierangelini, Alvin Wei Tian Ng, Nuno Raimundo, Ira Milosevic, Raimundo Freire, Yuliang Li, Sven Rottenberg, Dragomir B Krastev, Christopher J Lord, Madalena Tarsounas, Kristijan Ramadan

## Abstract

Poly (ADP-Ribose) Polymerase inhibitors (PARPi) induce cytotoxicity in homologous recombination repair (HRR)-deficient cancers by causing PARP1 to become trapped on chromatin, resulting in irreparable replication-associated DNA damage. Although increased clearance of trapped PARP1 from chromatin reduces the sensitivity of cancer cells to PARPi, details surrounding this process remain unclear. PARPi exposure is known to cause increased autophagy flux, whilst autophagy inhibition can hypersensitise cells to PARPi. Using various biochemical, cell biological and live imaging-based assays, we found that trapped PARP1 is cleared by nucleophagy, the selective autophagy of nuclear substrates. Specifically, the nucleophagy of trapped PARP1 was orchestrated by the selective autophagy receptor TEX264 and its partner segregase p97/VCP. TEX264 mediates this process by directly interacting with trapped PARP1, thus bridging PARP1 to the autophagosomal resident protein LC3 for processing via autophagy. Impeding this process, either chemically or genetically, heightened PARP1 trapping, leading to accumulation of protein aggregates, replication-associated DNA damage and cell lethality, re-sensitising PARPi-resistant cells to various PARPi. In conclusion, we show that nucleophagy acts in a cytoprotective manner to directly target PARPi-induced trapped PARP1 for degradation.

## Introduction

Since clinical trials began 20 years ago^1^, six Poly(ADP-Ribose) Polymerase inhibitors (PARPi) have been approved for the treatment of homologous recombination repair (HRR)-deficient breast, prostate, pancreatic and ovarian cancers^2–4^. This represents the first synthetic lethal targeted therapy, whereby HRR-defective cells, most commonly those with mutations in BRCA1 or BRCA2, display up to 1000-fold the selectivity to PARPi as their wild-type (WT) counterparts^5–7^. PARPi target the key DNA damage repair (DDR) enzyme PARP1, most notably involved in single-strand break (SSB) response^8^, replication fork stability^9^ and ligation of Okazaki fragments^10^.

All clinically approved PARPi are nicotinamide analogues which bind the catalytic site of PARP1^7^. Synthetic lethality was first attributed to catalytic inhibition of PARP1, an event that leads to the accumulation of SSBs, which in turn collapse into irreparable double-strand breaks (DSBs) during S-phase in HRR-defective cancers^5–7^. However, with the discovery that cytotoxicity of different PARPi does not directly correlate with their inhibitory effects and loss of PARP1 confers resistance (even in BRCA1 mutant cancer cells^11^), it was shown that PARP1 becomes tightly bound to DNA upon PARP inhibition, a phenomenon known as PARP1 trapping^12, 13^. The trapping potency of each PARPi strongly correlates with its cytotoxicity, demonstrating the importance of this mechanism for PARPi response^13–15^. Talazoparib is the most potent clinically approved PARP1 trapper^7, 14, 16^, used as a monotherapy in the treatment of human epidermal growth factor receptor 2 (HER2)-negative, BRCA-mutated locally advanced or metastatic breast cancer, and in clinical trials for various other cancers^3^. Trapping of PARP1 on endogenous DNA damage sites and unligated Okazaki fragments is thought to cause collisions with the replisome and fork stalling, which, in HRR/BRCA1/2-defective cells, collapse into DSBs that go unrepaired, leading to cell death^4, 12–14^.

Despite their early promise, PARPi resistance is a major concern in the clinic, with both acquired and *de novo* resistance common^8, 17, 18^. In ovarian cancer, 40% of patients with a germline BRCA mutation show no response to PARPi, either olaparib^18–20^ or Rucaparib^4, 21^. This highlights a lack of response to predictive biomarkers, with deleterious mutations in BRCA/HRR genes not always indicative of real-time HRR efficacy, degree of sensitivity to therapy or rate of resistance development^4, 22–24^. The only resistance mechanism broadly clinically validated is BRCA/HRR gene reversions and epigenetic regulation, which restore HRR, as observed in hundreds of patients^8, 25–27^. Pre-clinical evidence also points to (i) non reversion mutations that restore HRR in BRCA1-defective cancers through loss of DNA end resection inhibitors^28–31^, (ii) restored fork stability by inhibiting nuclease recruitment^32^, (iii) loss of poly(ADP-ribose) glycohydrolase expression^33^, (iv) upregulation of ABCB1 efflux transporter to prevent accumulation of PARPi^34–36^, and (v) PARP1 mutations that abolish trapping^11, 37^. Greater insight into trapped PARP1 biology is necessary to further understand clinically relevant resistance mechanisms, essential for identifying predictive biomarkers of resistance and improving treatment prospects.

One area of interest is the proteolytic processing of trapped PARP1. Both the metalloprotease SPRTN^38^ and the molecular unfoldase/segregase p97 (VCP in metazoans or Cdc48 in yeast)^39^ were implicated in extracting trapped PARP1 from chromatin. p97, via its co-factor UFD1, recognises trapped PARP1 SUMOylated and ubiquitylated by PIAS4 and RNF4, respectively, driving its extraction from chromatin via p97’s central pore^39^. p97 recently emerged as a promising druggable target for various cancers, and one of its inhibitors is in clinical trials^40^. p97 resolves various protein-induced DNA lesions on chromatin, such as TOP1ccs^41^ and trapped PARP1^39^, which can be covalently or non-covalently bound to DNA, respectively, causing cytotoxicity through replisome blockade^42^. Due to the pleiotropic roles of p97 in processes associated with chromatin, including DNA repair^43–48^, understanding both upstream regulation of p97 by its cofactors and downstream processing is key to developing this system as a target for therapy. Interestingly, our group recently uncovered the autophagy-dependent downstream processing of TOP1cc lesions cleared from chromatin by p97^49^. This pathway is reliant on the p97 cofactor and the selective autophagy receptor (SAR) TEX264. TEX264 bridges topoisomerase inhibitor-induced TOP1ccs to the autophagosome for degradation. This mechanism prevents the accumulation of cytotoxic protein aggregates, genome instability, and influences colorectal cancer patients’ response to the topoisomerase 1 inhibitor, Irinotecan.

Best characterised under starvation, bulk autophagy is a cellular homeostasis pathway whereby substrates become engulfed in elongating double-membrane phagophores, which form autophagosomes for fusion with hydrolase-containing lysosomes^50^. Autophagy of specific cargo, termed selective autophagy, can occur through association of cargo with ATG8-family proteins conjugated to the growing autophagophore membrane, such as lipidated LC3. This can occur through direct interaction between LC3 and cargo, but often requires SARs like TEX264^49, 51, 52^, which recognise specific substrates and bridge them to ATG8-family proteins^53, 54^. Many complete and ongoing clinical trials with autophagy inhibitors chloroquine and hydroxychloroquine, often as part of combination therapies, are showing positive results, but the development of other inhibitors is hindered by tolerability^55^. Combination therapies such as these form another key area of research in PARPi therapy^8^. All four FDA-approved PARPi have been shown to upregulate autophagy, giving a cytoprotective effect, in a multitude of cell lines and patient-derived xenografts^56–63^, suggesting it as a good target for combination therapies. However, it is essential to better understand the role of autophagy in PARPi response if this is to be successfully translated to the clinic.

Therefore, the search for predictive biomarkers and combination treatments to overcome PARPi resistance is of major clinical need. To further this goal, we investigated how trapped PARP1 is cleared and processed from chromatin. Through mass spectrometry, CRISPR screens and RNA-sequencing, we identified PARPi-induced upregulation of autophagic machinery as cytoprotective in response to PARPi and observed its interaction with trapped PARP1. Mechanistically, we demonstrated that autophagy plays a cytoprotective role through the clearance of trapped PARP1 by SAR TEX264 and its partner protein p97. Specifically, TEX264 binds directly to PARP1 and facilitates the removal of trapped PARP1 from chromatin by transporting it to the lysosome. The impairment of this pathway results in the accumulation of cytotoxic aggregates of PARP1 protein and genome instability, resulting in increased cell lethality, including in PARPi-resistant cells. Interestingly, we found that low TEX264 expression is positively correlated with improved overall survival in breast cancer patients whose tumours lack homologous recombination repair capacity. These clinical findings support the concept described here: that the TEX264-mediated nucleophagy pathway promotes the survival of HRR-deficient mammalian cells, specifically in BRCA1-deficient cases. Our results identify a TEX264-dependent nucleophagy pathway targeting trapped PARP1, which may have clinical relevance.

## Results

### Autophagy is upregulated upon PARPi treatment, with a cytoprotective effect

PARPi induces trapping of PARP1 on chromatin which is thought to be the major mechanism of PARPi cytotoxicity^12, 13, 15, 64, 65^. Recent work has suggested that trapped PARP1 can be extracted from chromatin by the enzymatic activities of the ATPase p97 and SPRTN protease, similar to other DNA-protein crosslinks (DPCs)^38, 39, 66, 67^, potentially contributing to PARPi resistance. Through an RNA-seq approach, we explored which protein homeostasis systems may be differentially expressed upon PARPi treatment, suggesting their potential involvement in trapped PARP1 repair. We performed RNA-seq on triple-negative breast cancer (TNBC) CAL51 cells after 24 hours of treatment with strong trapping PARPi talazoparib. As expected, we observed upregulation of genes related to DNA damage repair, replication stress, apoptosis, G1/S cell cycle checkpoint and mitotic checkpoint (Fig. S1A, Table S1). Top hits include CDKN1A and BTG2, involved in p53-dependent G1/S checkpoint signalling in response to DNA damage^68^, MDM2, a p53 regulator which also has p53-independent roles in the regulation of DNA synthesis and repair^69^ and BAX, a pro-apoptotic protein^70^ (Fig. 1A, Table S2). By probing with a comprehensive gene list for the autophagy pathway^71^, we observed upregulation of 31 autophagy-related genes upon treatment. Significantly upregulated genes included SESN1, SESN2 and DRAM1 (Fig. 1A, Table S2), which are well-established to be transcriptionally activated by p53 under genotoxic stress conditions to promote autophagy^72^. In accordance with this, numerous studies show upregulation of autophagy in cells and patient-derived xenografts treated with PARPi^56–63, 73^. We also probed with a GO term list for proteasome-mediated ubiquitin-dependent catabolic processes (GO:0043161). Only 15 genes were differentially expressed, 7 significantly upon treatment, and when we used a GO list specific for the proteasome complex itself (GO:0000502), we observed no differential expression. In fact, most proteasome-associated genes identified were E3 ligases, components of E3 ligases or E2 conjugating enzymes well-established in DNA damage response and cell cycle regulation in response to stress, such as MDM2, FBXW7 and UBE2C (Table S2). Core proteasome protein transcripts are some of the most consistently expressed in human cancer cells^74^. However, whilst their lack of differential expression is unsurprising, very few genes known to positively regulate the proteasome (5 genes from GO:1901800) were differentially expressed (Table S2). This implies that PARPi may not induce proteasomal degradation to the same extent as autophagy.

**Figure 1:**
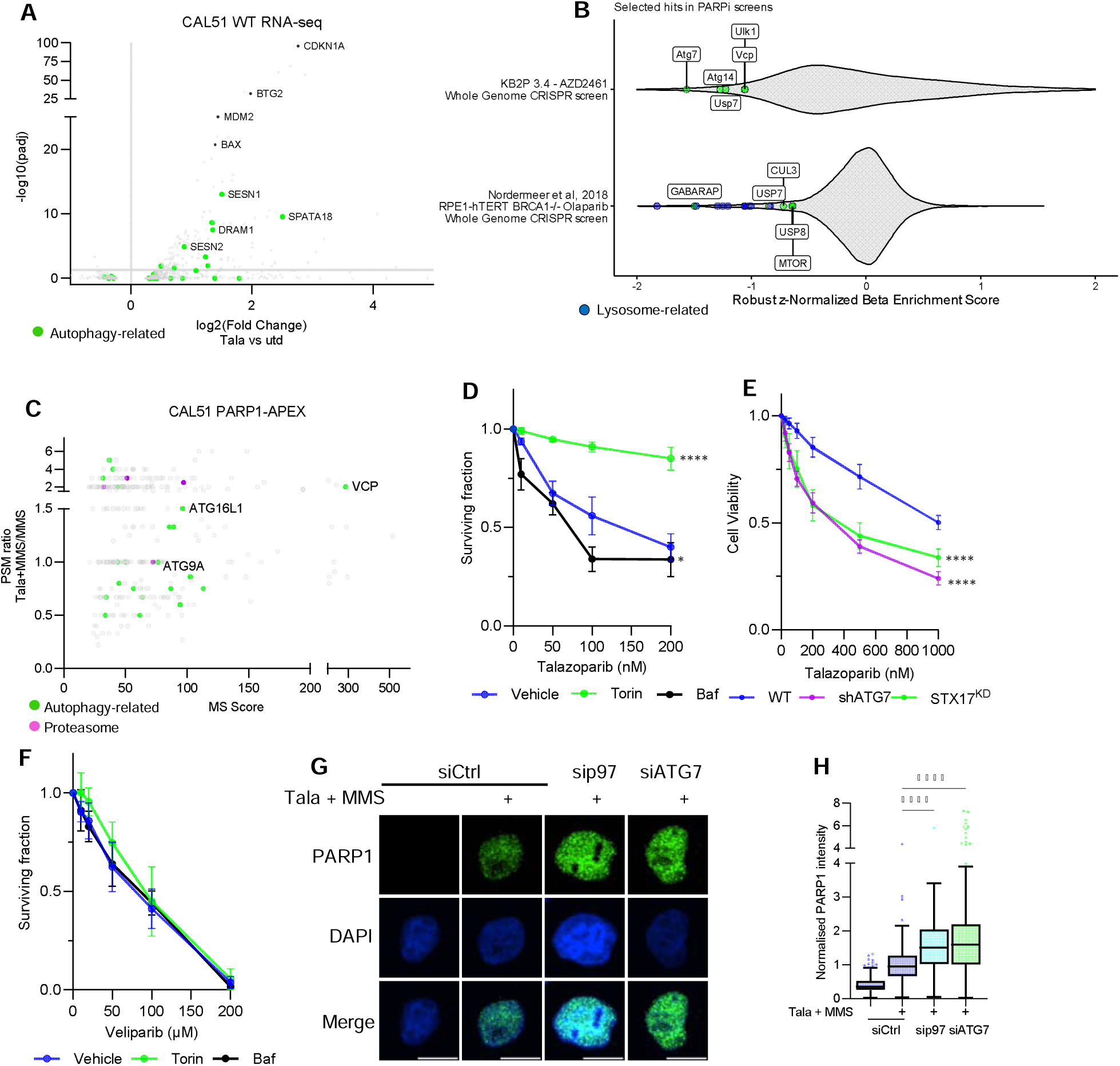
PARPi-induced upregulation of autophagy has a cytoprotective, trapping-related effect. **(A)** Volcano plot showing differential gene expression in CAL51 cells comparing talazoparib (100 nM, 24 hrs) to untreated by RNA-seq. The grey line indicates the adjusted p-value of 0.05. **(B)** Violin plots showing selected autophagy-related depletion hits from whole genome CRISPR screens performed in KB2P 3.4 (BRCA2 knockout) and RPE-hTERT BRCA1^-/-^ treated with AZD2461 and olaparib, respectively^31, 76^. **(C)** Mass spectrometry data from Krastev *et al.,* (2022)^39^ of PARP1 interactions that are enriched under PARP1-trapping conditions (Tala + MMS) by PARP1WT–Apex2–eGFP proximity labelling. **(D)** Colony formation assay in HeLa cells treated with talazoparib and bafilomycin A1 (25 nM) (Baf) or torin-1 (150 nM) for 24 hrs. Statistical analysis by 2-way ANOVA. **(E and F)** Cell viability measured by resazurin assay in HeLa cells, either WT, dox-inducible shATG7, or stable STX17^KD^, treated with **(E)** talazoparib or **(F)** veliparib for 24 hrs, followed by 48 hrs recovery. **(G)** Immunofluorescence with detergent pre-extraction to detect trapped PARP1 foci in cells depleted for the indicated proteins using siRNA and treated with talazoparib and MMS. Scale bar is 10 μm. **(H)** Quantification of (G) from >300 cells across 2 biological repeats, with mean +/- SEM shown and statistical analysis by one-way ANOVA. Normalisation is by dividing each point by the mean of siCtrl treated within each repeat.

To further explore whether autophagy is important in cellular response to PARPi, we explored two previously published whole genome CRISPR screens for loss of autophagy factors affecting sensitivity to PARPi. One of the screens was performed in BRCA2^-/-^ TRP53^-/-^ mouse mammary tumour cell line KB2P-3.4^75^ treated with PARPi AZD-2461^76^, while the other was performed in RPE-hTERT TP53^-/-^ BRCA1^-/-^and treated with olaparib^31^. Following functional enrichment analysis across four different databases, we identified several significantly represented gene sets relating to autophagy amongst the sensitivity candidates (Fig. S1B), suggesting that loss of some genes belonging to these terms caused sensitivity to PARPi. Notably, “regulation of Macroautophagy” emerged as one of the most significantly enriched pathways. Specifically, we found that the loss of a handful of core autophagy factors significantly sensitises cells to PARPi. These include ATG7, ATG4, ULK1 and GABARAP. Loss of regulators of autophagy USP7^77^, USP8^78, 79^, CUL3^80^ and p97/VCP^81^ are also implicated in cell sensitivity to PARPi (Fig. 1B). Moreover, several of the depleted genes are involved in lysosome biogenesis and function, or are subunits of v-ATPase, which is key in autophagic degradation (Fig. 1B, S1C). From these data, impaired autophagy seems to cause increased sensitivity to two different PARPi and in cell lines with either BRCA1 or BRCA2 deficiency, which constitute the majority of tumours treated with PARP inhibitors in the clinic.

To further understand the role of autophagy in PARPi response, we turned to mass spectrometry data from our previous work^39^, that identified trapped PARP1-interacting proteins defined by PARP1-APEX proximity labelling. Interestingly, in gene set enrichment analysis, autophagy appeared as the 8^th^ most significant gene set enriched under PARP1 trapping conditions (Fig. S1D). To explore this, we probed this data set with the same autophagy gene list used when analysing RNA-seq data and identified 22 autophagy-related proteins interacting with trapped PARP1 (Fig. 1C). Alongside many autophagy regulators, we identified ATG9A and ATG16L1, two key regulators of autophagosome biogenesis, indicating the proximity of trapped PARP1 with the core autophagy machinery. Despite not being enriched in our RNA-seq data, the proteasome ranked 14^th^ by significance in gene set enrichment analysis (Fig. S1D), with 4 proteasome subunits (PSMA6, PSMB5, PSMD2 and PSMD12) identified in the PARP1 interactome (Fig. 1C). Proteasomal degradation of trapped PARP1 in basal conditions is well-characterised^82–84^, and this indicates that it may also degrade trapped PARP1. The presence of so many core autophagy genes in the trapped PARP1 proteome implies a potential direct role of autophagy in regulating chromatin-bound PARP1. When considering this alongside its upregulation by RNA-seq (Fig. 1A) and in previous literature^57^, as well as the PARPi-sensitising effect that loss of autophagy genes caused in CRISPR screens (Fig. 1B, S1B), autophagy seems an attractive target for further exploration.

We began by exploring how modulation of autophagy impacts cellular response to PARPi. We treated cells with talazoparib in combination with either bafilomycin A1, to impede autophagy through inhibition of lysosome acidification, or with torin-1, to boost autophagic flux through mTOR inhibition. Torin-1 treatment caused a striking resistance to talazoparib, whilst inhibition of autophagy with bafilomycin A1 increased sensitivity (Fig. 1D). As mTOR has multiple functions, we combined torin-1 treatment with depletion of ATG7, an E1-like enzyme involved in conjugating LC3 to the autophagosome membrane during phagophore formation and expansion^50, 55^. This confirmed that the PARPi resistance induced by torin-1 is due to increased autophagy flux, as resistance to talazoparib was partially but significantly reversed by ATG7 depletion (Fig. S2A). We also confirmed the association of autophagy and PARPi genetically through inhibition of autophagy either by depletion of ATG7 or syntaxin-17, an autophagosomal SNARE protein that regulates multiple autophagic processes, including autophagosome membrane fusion with the lysosome^85, 86^. Depletion of either of these autophagy factors increased the sensitivity of cells treated with talazoparib (Fig. 1E, S2B). Depletion of ATG9A, a lipid scramblase involved in autophagophore formation and identified as an interactor of PARP1 (Fig. 1C), also caused increased sensitivity to talazoparib (Fig. S2C, S2D). This is in line with our findings from CRISPR screen data (Fig. 1B, S1B) and various recent studies showing the effect of autophagy inhibition in increasing sensitivity to PARPi^56, 57, 59–62^. This suggests that, as seen before, upregulation of autophagy upon PARPi treatment has a cytoprotective effect.

The finding that PARPi sensitivity is heightened by autophagy impairment has been demonstrated before, and various explanations have been tendered for the importance of autophagy upregulation upon PARPi treatment. This includes PARPi- induced upregulation of PTEN to promote cytoprotective autophagy in response to PARPi-induced ROS^62^ and nuclear localisation of p62, indirectly causing upregulated homologous recombination^87^. In any of the conditions which induced either sensitivity or resistance to PARPi, we observed no effect on the nuclear localisation of p62 (Fig. S2E), implying an alternative role of autophagy.

Remarkably, we did not observe the same cytoprotective effect of autophagy when cells were treated with veliparib, a PARPi that inhibits PARP1 catalytic activity but causes no trapping^14, 16^, with neither bafilomycin A1 or torin-1 treatment affecting cell sensitivity (Fig. 1F). This implies that the observed effect is linked to PARP1 trapping. Using detergent pre-extraction to remove soluble proteins, but not those that are tightly bound to chromatin, we visualised trapped PARP1 on chromatin by immunofluorescence under trapping conditions, when cells were treated with talazoparib and a low dose of the alkylating agent MMS^39^ (Fig. 1G and H, S2F). PARP1 is only visualised on chromatin under trapping conditions, confirming the efficacy of this assay for detecting specifically trapped PARP1. Trapped PARP1 levels were significantly increased (∼1.7-fold) upon autophagy inhibition by depletion of ATG7, a similar increase as was seen in p97-depleted cells, which are known to accumulate trapped PARP1^39^. Altogether, this implies that autophagy upregulation upon PARPi treatment has a cytoprotective effect by restricting accumulated trapped PARP1. Whilst an association between PARPi sensitivity and autophagy flux has previously been shown in multiple studies, we believe this is the first to show that autophagy plays a direct role in modulating the PARPi response.

### Trapped PARP1 is processed by autophagy

As trapped PARP1 levels were increased upon autophagy inhibition and early-stage autophagy core machinery was found in proximity to trapped PARP1 by mass spectrometry (Fig. 1), we posited that trapped PARP1 could be cleared by autophagy. To explore this, intact lysosomes were isolated from HeLa and CAL51 cells expressing lysosomal transmembrane protein TMEM192-3HA by a method known as LysoIP to analyse lysosomal contents (Fig. 2A)^88^ in control and PARP1 trapping conditions. Remarkably, PARP1 was localised in the lysosome and accumulated considerably under PARP1 trapping conditions in both HeLa and CAL51 cells (Fig. 2B, 2C, S3A, S3B). Lysosomal PARP1 levels upon trapping were increased further by treatment with Bafilomycin A1, which neutralises lysosome pH, stabilising its contents, further confirming that PARP1 localises in the lysosome (Fig. 2B, 2C). PARP1 was more extensively localised to the lysosome when treated with the strong PARP trapping inhibitors talazoparib and niraparib, compared to veliparib, a weaker trapper of PARP1 (Fig. 2D and E, S3C and D). In CAL51 cells expressing either WT PARP1 (PARP1^WT^) or a trapping-impaired mutant PARP1^del.p.119K120S^, only PARP1^WT^ was localised to the lysosome (Fig. S3A, S3B). Whole cell levels of PARP1^del.p.119K120S^ were lower than PARP1^WT^, but even when its lysosomal level was normalised against this across 7 biological repeats, we observed a robust reduction in lysosomal PARP1^del.p.119K120S^ compared to PARP1^WT^ (Fig. S3B), highlighting that only trapped PARP1 is processed in this way. To further validate this, we used a modified version of the well-recognised mCherry-GFP autophagy reporter assay, whereby cells were transfected with a PARP1 construct tagged with mCherry and GFP (Fig. S3E). In most cellular compartments, including the nucleus, both mCherry and GFP will fluoresce and co-localise. However, in the acidic environment of the lysosome, GFP is quenched, so only the mCherry signal is observed^89^ (Fig. 2F). PARP1 is predominantly localised in the nucleus, where the red and green signals were observed most strongly and co-localise. However, under trapping conditions, cytosolic red puncta of PARP1 formed (Fig. 2G and H), indicating its localisation to the lysosome. These fluoresced both red and green when bafilomycin-A1 was added to neutralise the lysosomal pH and prevent quenching of the GFP signal, confirming the specificity of this assay for lysosome-localised PARP1 (Fig. 2G and H). Depletion of ATG7 reduces the number of cytosolic puncta to background levels seen without treatment, further confirming that PARP1 exits the nucleus in an autophagy-dependent manner upon trapping. These cytosolic puncta are no longer observed. To further confirm the localisation of PARP1 to the lysosome after treatment, we visualised mCherry-GFP-tagged PARP1 in cells stained with a lysosome dye by live imaging (Fig. 2I). We observed puncta emerging from the nucleus within ∼5 minutes of talazoparib and MMS treatment (treatment added at 2:30), with multiple puncta appearing across the 2-hour assay (Movie 1). These puncta localised with lysosomes, for around 20 minutes until they had been degraded and lysosomes dispersed (Fig. 2I: best view at 9min in the lower panel; PARP/red signal engulfed by lysosome, and Movie 1). Using rendering, we were able to better distinguish the green signal from background noise and visualise lysosomes with better resolution. Remarkably, we observed lysosomes moving close to the nuclear periphery within minutes of treatment and strongly co-localising with PARP1 signal as it exits the nucleus (Fig 2I and Movie 2). Together, this further confirms that PARP1 localises to the lysosome under PARP trapping conditions.

**Figure 2:**
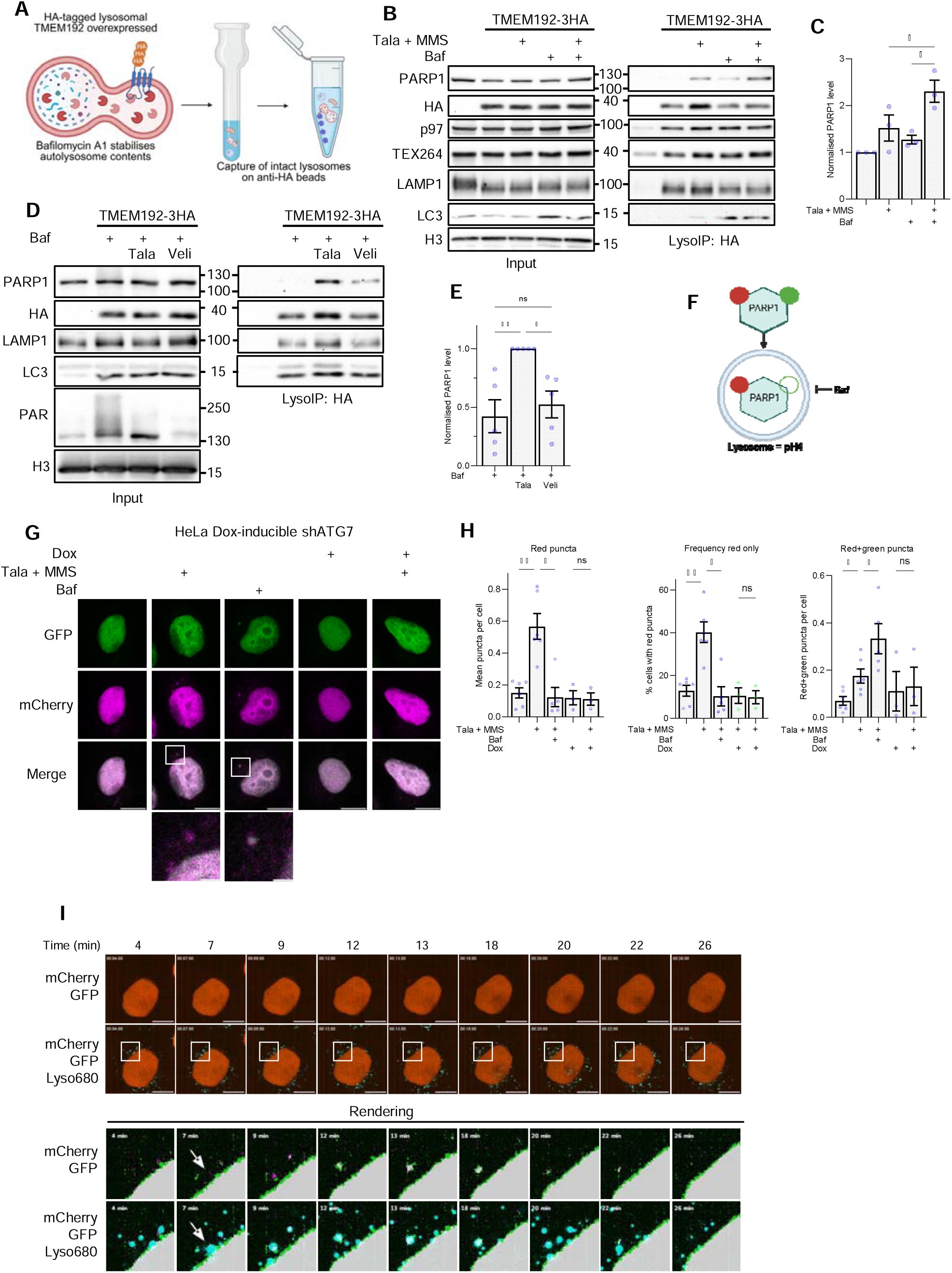
Trapped PARP1 is processed by selective autophagy. **(A)** Schematic showing methodology of LysoIP, whereby TMEM192-3HA is stably expressed in cells treated with bafilomycin A1, then immunoprecipitated to isolate intact lysosomes. **(B)** LysoIP in HeLa cells treated with the indicated drugs, showing the accumulation of PARP1 in the lysosome under trapping conditions. **(C)** Quantification of PARP1 levels in (B) from 3 biological repeats, normalised to untreated. **(D)** LysoIP in HeLa cells treated with either talazoparib (200 nM) or veliparib (5 µM) for 3 hrs. **(E)** Quantification of PARP1 levels in (D) from 3 biological repeats, normalised to talazoparib-treated. **(F)** Schematic showing the methodology of the mCherry-GFP reporter assay. PARP1 tagged with mCherry and GFP is transiently expressed in cells and visualised by either fixed or live imaging. GFP is quenched in acidic environments, so only the red signal is detected in the lysosome. This can be reversed by treatment with bafilomycin A1, which neutralises the lysosome. **(G)** Fixed images from the mCherry-PARP1-GFP reporter assay in HeLa dox-inducible shATG7 cells treated with the indicated drugs for 3 hrs. Images have a scale bar of 10 µm. Zoom images show puncta indicated by white boxes with a scale bar of 2 µm. **(H)** Quantification of mCherry-GFP puncta in (G) from >3 biological repeats for each condition with statistical analysis by one-way ANOVA. **(I)** Snapshots from live cell imaging of HeLa cells transfected with mCherry-PARP1-GFP and stained with LysoView 680. Time stamps indicate time since the beginning of the assay. Talazoparib and MMS were added at 2:30. Scale bar is 5 µm. Bottom panels show images processed by rendering using the TrackMate plugin on ImageJ.

To confirm whether this lysosomal localisation of trapped PARP1 is dependent on autophagy, we depleted ATG7 and assessed lysosomal PARP1 levels by both lysoIP and the mCherry-PARP1-GFP reporter assay. Depletion of ATG7 almost entirely abolished both the recruitment of PARP1 to the lysosome under trapping conditions (Fig. 3A, 3B) and the accumulation of red PARP1 puncta (Fig. 2G, H). We observed the same effect when autophagy was inhibited through depletion of syntaxin-17 (Fig. 3C and D), ATG9A or Beclin-1 (Fig. 3E and F). As these proteins have different functions in varying stages of autophagy, this comprehensively demonstrates that PARP1 is processed by autophagy under trapping conditions.

**Figure 3:**
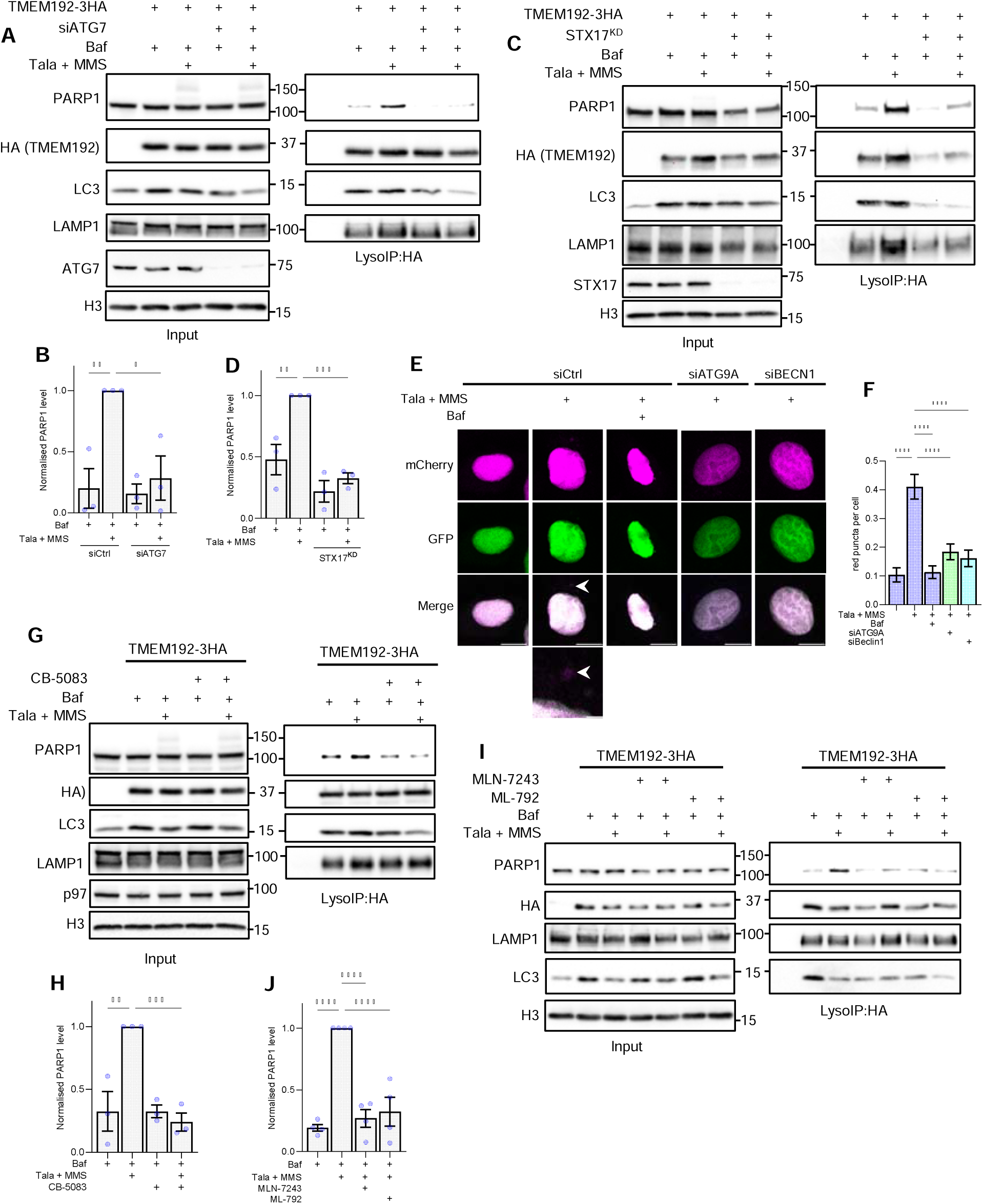
Trapped PARP1 is processed by p97, ubiquitin and SUMO-dependent autophagy. **(A)** LysoIP in HeLa cells with depletion of ATG7 by siRNA. **(B)** Quantification of PARP1 levels in (A), normalised to the treated control. **(C and D)** As in (A and B), but in either WT cells or cells with stable syntaxin-17 knockdown (STX^KD^). **(E)** Fixed images from the mCherry-PARP1-GFP reporter assay in HeLa cells depleted of either ATG9A or Beclin-1 and treated with the indicated drugs for 3 hrs. Images have a scale bar of 10 µm. Zoom images show puncta indicated by white boxes with a scale bar of 2 µm. **(F)** Quantification of mCherry-GFP puncta in (E). **(G)** LysoIP in HeLa cells treated with talazoparib, and MMS combined with p97i CB-5083 (10 μM). **(H)** Quantification of PARP1 levels in (G), normalised to the treated control. **(I)** LysoIP in HeLa cells treated with the indicated drugs, including ubiquitination inhibitor MLN-7243 (5 μM) or SUMOylation inhibitor ML-792 (1 μM). **(J)** Quantification of PARP1 levels in (I), normalised to the treated control. All graphs present at least 3 biological repeats with statistical analysis by one-way ANOVA.

### Autophagosomal processing of trapped PARP1 is dependent on p97, ubiquitination and SUMOylation

As we have demonstrated the processing of trapped PARP1 by selective autophagy, we next wanted to explore if this occurs downstream of the previously described p97-dependent extraction of trapped PARP1 from chromatin^39^. Importantly, besides its chromatin role in removing various DNA-associated substrates, p97 was also implicated in autophagy and moves between the cytoplasm and nucleus depending on cellular need ^90–93^. However, the role of p97 in autophagy remains to be fully understood. Using lysoIP to assess PARP1 levels in the lysosome under trapping conditions, we inhibited p97 with the specific and clinically relevant inhibitor CB-5083^40^. CB-5083 treatment significantly reduced the accumulation of PARP1 in the lysosome (Fig. 3G and H), suggesting a role of p97 in mediating this process. This is in accordance with the impaired removal of PARP1 from chromatin when p97 is inhibited, as observed previously ^39^.

p97 is recruited to trapped PARP1 partly through SUMOylation and subsequent ubiquitylation^39^, with the latter often involved in selective autophagy. Inhibition of either ubiquitination or SUMOylation caused reduced lysosomal engulfment of PARP1 under trapping conditions (Fig. 3I and J). Trapped PARP1 ubiquitination has previously been shown to be mediated by the SUMO-targeted E3 ubiquitin ligase RNF4^39^. Modulation of RNF4 activity, either by expression of its dominant negative enzymatic inactive M136A+R177A variant (RNF4^DN^) or over-expression of its WT (RNF4^WT^), did not affect lysosomal PARP1 levels (Fig. S3F and G). This negates the role of RNF4-mediated ubiquitination of PARP1 in regulating trapped PARP1 processing by autophagy. Instead, RNF4-mediated ubiquitination of trapped PARP1 likely mediates its proteasomal degradation, as has been described for most of its other substrates^94, 95^, including PARP1 in response to heat shock^96^. In the p97-RNF4 pathway previously described, p97 is recruited to ubiquitinated PARP1 by the co-factor UFD1^39^. As with RNF4 modulation, depletion of UFD1 did not affect trapped PARP1 levels in the lysosome (Fig. S3H and I). Altogether, this suggests that the previously described RNF4-UFD1-p97 pathway acts separately from the processing of trapped PARP1 by autophagy. p97 is still involved but must rely on other unknown SUMO or ubiquitin ligases and p97 co-factor(s) acting to facilitate this pathway.

### Selective autophagy of trapped PARP1 is TEX264-dependent but nuclear pore-independent

We have described the processing of trapped PARP1 in the lysosome and shown that this occurs by autophagy in a p97-dependent manner. Autophagy can selectively target substrates through selective autophagy receptors (SARs), which bridge specific, often ubiquitinated, cargo to ATG8 family proteins (including LC3) in a growing autophagosome^54, 97^. To further understand the processing of trapped PARP1, we wanted to identify the SAR responsible for its processing. Interestingly, TEX264 has recently been identified as a SAR for the endoplasmic reticulum under starvation conditions^51, 52, 98, 99^, and as a p97 co-factor and SAR essential for the removal of TOP1cc from chromatin by autophagy^41, 49^. Due to the similarity between TOP1cc and trapped PARP1 in blocking replication machinery, alongside the ability of TEX264 to act as both a SAR and p97 co-factor^41,49, 98^, we explored whether TEX264 may act in this pathway. We tested TEX264^-/-^ cells for their sensitivity to talazoparib and niraparib and found that loss of TEX264 caused a profound increase in sensitivity of both HeLa and CAL51 cells (Fig. 4A-D, S4A). Interestingly, and similarly to modulation of autophagy (Fig. 1D, 1F), the same effect was not observed in cells treated with reduced-trapping PARPi veliparib (Fig. 4E and F). This suggests that the function of TEX264 in protection from PARPi-induced cell death is linked to trapped PARP1. To confirm this, TEX264 was depleted and PARPi sensitivity assessed in cells expressing either PARP1^WT^ or a mutant previously shown to induce PARPi resistance by preventing PARP-trapping (p.43delIM;44F>I)^11^. TEX264 depletion only hypersensitised PARP1^WT^ cells to talazoparib and not the cells expressing the trapping-defective mutant (Fig. 4G). A considerable accumulation of trapped PARP1 was also observed after either depletion or knockout of TEX264 in three different human cell lines, including the HeLa cervical carcinoma cell line and two TNBC cell lines, CAL51 and MDA-MB231 (Fig. 4H-J, S4B-F). The level of trapped PARP1 observed was equivalent to the depletion of a previously identified cofactor UFD1, supporting that TEX264 acts as a physiologically relevant modulator of trapped PARP1 levels.

**Figure 4:**
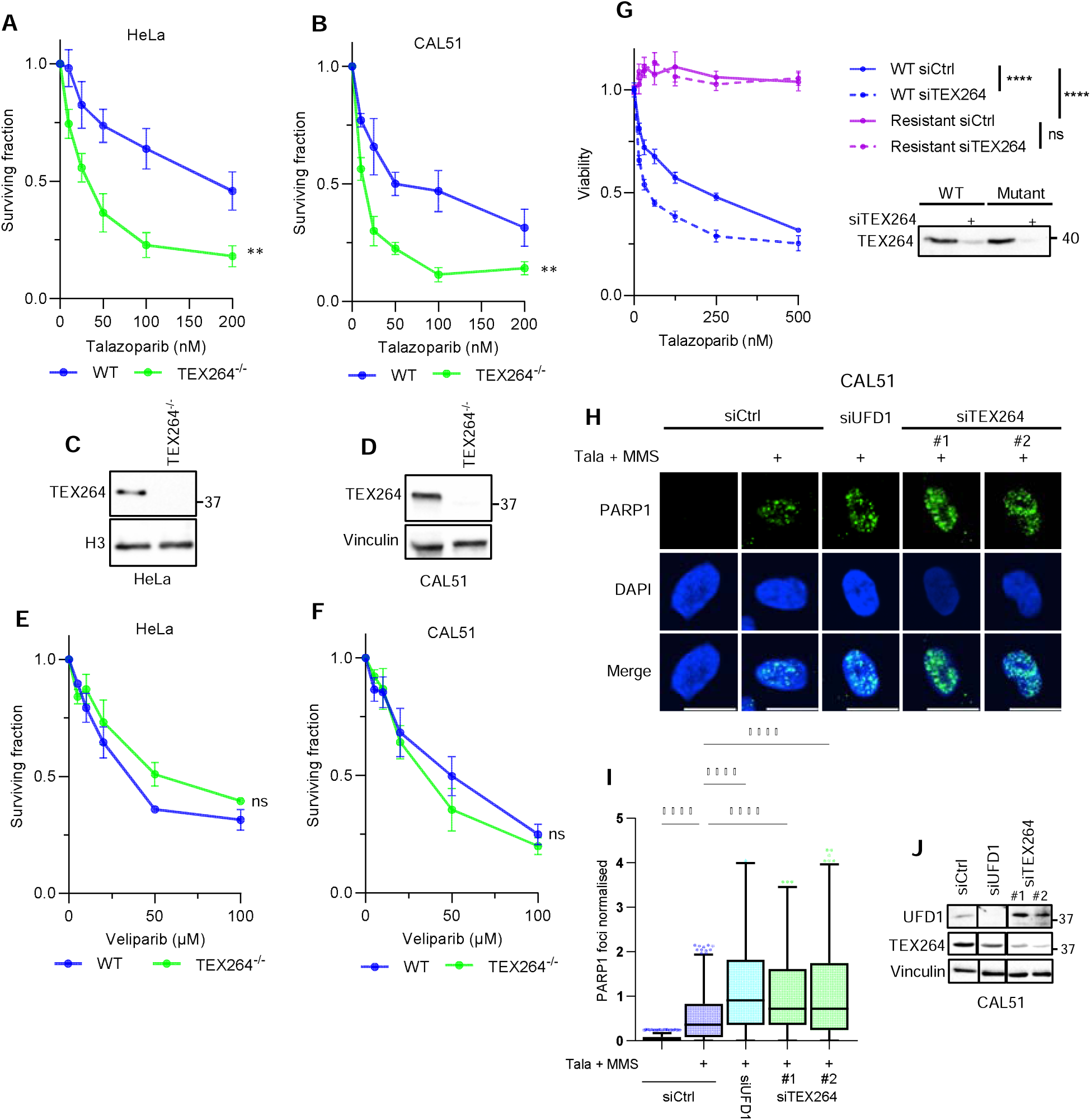
TEX264 acts as a regulator of PARP1 trapping. (A and. **B)** Colony formation assay in HeLa (A) and CAL51 (B) WT and TEX264^-/-^ cells treated with talazoparib for 24 hrs. Statistical analysis by two-way ANOVA. **(C and D)** Immunoblotting validating knockout of TEX264 in HeLa (C) and CAL51 (D). **(E and F)** As in (A) and (B) but with veliparib treatment instead of talazoparib. **(G)** Cell viability by resazurin assay in HeLa cells expressing either PARP1^WT^ or DNA-binding mutant PARP1^(p.43delIM;44F>I),^ which are resistant to PARPi. Cells were depleted of TEX264 with siRNA and treated with talazoparib for 24 hrs, followed by 48 hrs of recovery. Statistical analysis by two-way ANOVA. Immunoblot validates PARP1 expression and TEX264 depletion. **(H)** Images of trapped PARP1 foci visualised by immunofluorescence with detergent pre-extraction after talazoparib and MMS treatment in CAL51 cells depleted of either UFD1 or TEX264 with siRNA. Scale bar = 10 µm. **(I)** Quantification of PARP1 foci in (H) from 3 biological repeats with statistical analysis by one-way ANOVA. **(J)** Immunoblotting validating depletion of UFD1 and TEX264 for (H and I).

To further understand the role of TEX264 in this pathway, we turned to previously published TEX264 interactomes. In two such interactomes, PARP1 is identified as one of the top hits^41, 51^, indicating an interaction between these proteins. In line with this, TEX264 was co-immunoprecipitated from chromatin-bound GFP-tagged PARP1 (Fig. 5A). As expected from previous work^39^, p97 interacted with PARP1 specifically under trapping conditions (Fig. 5A). The interaction between TEX264 and chromatin-bound PARP1 also increased ∼40% after treatment (Fig. 5A and B). Interestingly, depletion of TEX264 caused a reduction in the level of p97, both localised to chromatin and to trapped PARP1, similar to what has previously been observed with UFD1 depletion (Fig. 5A and C)^39^. This implies that TEX264 acts as a p97 cofactor to aid in its recruitment to trapped PARP1. To determine whether PARP1 and TEX264 interact directly, we expressed and purified GFP-PARP1 and TEX264 from the *E.coli* expression system, and performed an in vitro pull-down assay by isolating PARP1 over PARP1-trap beads. Our results demonstrated that TEX264 directly binds to PARP1, but not to the PARP1 trap beads alone (Figure S5A). To explore this interaction further, we performed hydrogen-deuterium eXchange mass spectrometry, where a difference in deuterium uptake in TEX264 peptides indicates a region of interaction with PARP1. Multiple regions showed a strong difference in deuterium uptake, most predominantly in an α-helix immediately following the gyrase inhibitory-like domain (GyrI-like), and from S272 to the C-terminus, where other motifs such as the SHP and LIR are located (Fig 5D and S5B). A further *in vitro* pull-down using either a fragment of TEX264 containing only the GyrI-like domain or the C-terminal half, containing both regions of interest, demonstrated that only the C-terminal half interacts with TEX264 (Fig S5C). We tested the importance of both these regions for TEX264-PARP1 interaction in cells by expressing the TEX264 variants with these regions deleted (TEX264-272 and TEX264- Δα-Helix; Fig. 5E and S5D). Whilst loss of the α-helix had no impact on the interaction (Fig S5D), loss of the C-terminus from S272 onwards completely abolished interaction of TEX264 with chromatin-bound PARP1 (Fig 5E). This confirmed that a motif in the extreme C-terminus is required for PARP1-TEX264 interaction.

**Figure 5:**
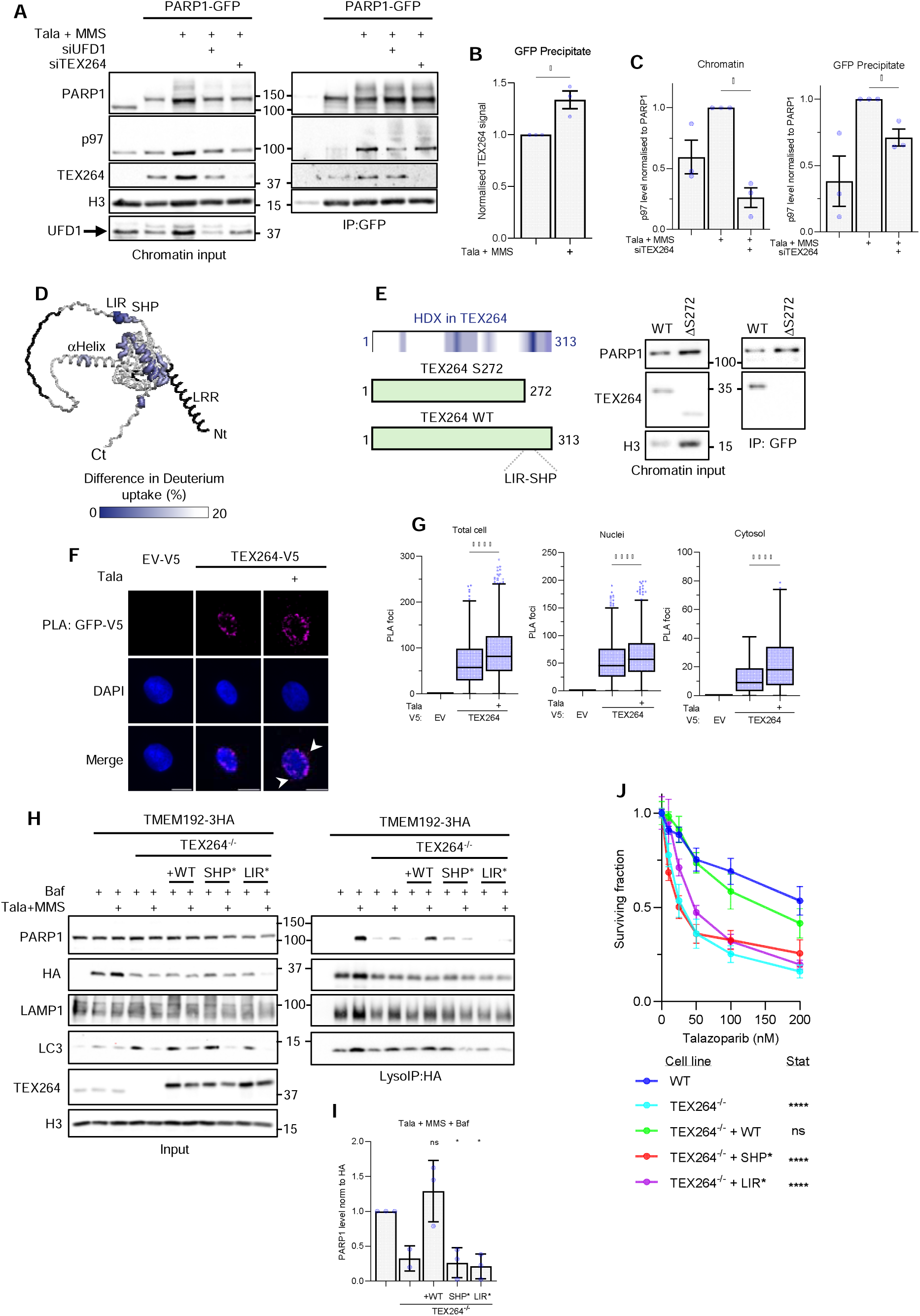
TEX264 serves as a p97-mediated selective autophagy receptor for trapped PARP1. **(A)** Co-immunoprecipitation of GFP from the chromatin fraction of CAL51 cells stably expressing PARP1-GFP after treatment with talazoparib and MMS for 3 hrs. Cells were depleted of the indicated genes using siRNA. **(B)** Quantification of TEX264 levels in the GFP-precipitate fraction is shown in (A) from 3 biological repeats 3 repeats with statistical analysis by students’ unpaired t-test. The TEX264 signal is normalised to the PARP1 signal to account for differences in binding to beads and to the untreated level. **(C)** Quantification of p97 levels in the chromatin fraction (left) and GFP-PARP1-co-IP (right), shown in (A) from 3 biological repeats 3 repeats with statistical analysis by students’ unpaired t-test. p97 levels were normalised by dividing by the PARP1 levels. **(D)** The structure of TEX264 with enlarged blue regions indicating areas of altered deuterium uptake when combined with PARP1 in hydrogen-deuterium eXchange mass spectrometry (HDX-MS) with 30 sec of labelling. The α-helix and C-terminal regions are labelled as areas with altered deuterium uptake, indicating that they interact directly with PARP1. The heatmap indicates a difference in deuterium uptake from 0-20%. Nt and Ct indicate the N- and C-terminus, respectively. **(E)** Co-immunoprecipitation of GFP from the chromatin fraction of HeLa cells stably expressing PARP1-GFP after treatment with talazoparib and MMS for 3 hrs. Cells expressed either TEX264^WT^ or a variant lacking the C-terminal region beyond S272. Schematics (left) show the TEX264 variants compared to a map showing sites of interest highlighted by HDX-MS. **(F)** Proximity ligation assay (PLA) between GFP and V5 in cells stably expressing PARP1-GFP and either TEX264-V5 or empty vector (EV)-V5 after treatment with talazoparib for 3 hrs. Scale bar is 10 µm in the top panels. Bottom panel shows a zoom of the yellow box with a scale bar of 5 µm. **(G)** Quantification of foci in (F) in whole cell, cytosol or nuclei from 3 biological repeats as shown by Tukey box plot with statistical analysis by one-way ANOVA. **(H)** LysoIP in HeLa cells stably expressing TMEM192-3HA with or without TEX264^-/-^. TEX264-V5 variants are transiently expressed where indicated. **(I)** Quantification of PARP1 levels in (H) from 3 biological repeats, except for TEX264^-/-,^ which was included in 2 repeats, normalised to the WT control. Statistical analysis by one-way ANOVA compared to WT. **(J)** Colony formation assay in CAL51 WT, TEX264^-/-^ or TEX264^-/-^ cells stably expressing indicated TEX264 variants, treated with talazoparib for 24 hrs. 6 technical replicates from 2 biological repeats with statistical analysis by two-way ANOVA compared to WT.

To visualise TEX264-PARP1 interaction beyond the chromatin context, we carried out proximity ligation assays (PLA) between GFP-tagged PARP1 and V5-tagged TEX264 (Fig. 5F, S5E). We were able to visualise this interaction and, as with our co-IP experiments, observed that it increased in the nucleus (Fig. 5F and G). PARP1-TEX264 PLA signal is partially localised around the nuclear periphery, where TEX264 is known to localise due to its transmembrane N-terminal leucine-rich region^41^. Surprisingly, despite the predominantly nuclear localisation of PARP1^100^, we observed PLA signal increasing considerably after PARPi treatment in the cytosol (Fig. 5F and G). This cytosolic PARP1-TEX264 interaction further validates our hypothesis that TEX264 may act as a SAR of trapped PARP1.

The function of TEX264 in selective autophagy and as a p97 co-factor is reliant on its LC3-interacting region (LIR)^51, 52^ and p97-interacting SHP motif^41, 101^, respectively. To explore its role in the autophagosomal processing of trapped PARP1, we performed lysoIP in TEX264^-/-^ cells (Fig. 5H). Interestingly, the loss of TEX264 impaired the accumulation of PARP1 in the lysosome under trapping conditions. Complementation of the TEX264-null background with TEX264^WT^ restored lysosomal PARP1 levels. However, mutation of the LIR domain ablated lysosomal PARP1 (Fig. 5H and I). As the LIR is a crucial motif for bridging substrates to LC3 localised in the autophagosomal membrane^97^, this suggests that TEX264 acts as a SAR for trapped PARP1. Similar to p97 inhibition, mutation of the TEX264 SHP domain also impaired PARP1 accumulation in the lysosome under trapping conditions (Fig. 5H and I). Remarkably, the TEX264^SHP*^ and TEX264^LIR*^ variants were also unable to rescue PARPi sensitivity in TEX264^-/-^ cells (Fig. 5J, S5F), further confirming the importance of these two functions of TEX264 in this pathway.

As part of its function as a SAR, TEX264 localises to the ER and both the inner and outer nuclear membranes, as has been shown with electron microscopy^102^. We confirmed this using PLA between laminA/C-GFP and TEX264-V5 (Fig. S6A). However, how PARP1 is transported out of the nucleus for autophagosomal processing remains unclear. Lamin-A/C, another substrate of nucleophagy ^103, 104^, is phosphorylated by ATR in response to DNA damage, promoting local nuclear envelope rupture^105, 106^. Recent work into TEX264-dependent autophagosomal processing of TOP1cc demonstrated that lysosomal localisation of TOP1 is ATR-dependent but not reliant on nuclear pore activity^49^. To assess if the same is true for trapped PARP1 processing, lysoIP was carried out under trapping conditions in cells treated with either the ATR inhibitor, VE-822, or the nuclear pore inhibitor, leptomycin B (Fig. S6B-E). In line with previous work, leptomycin B did not affect lysosomal PARP1 levels (Fig. S6B and C) while ATRi caused a significant reduction (Fig. S6D and E). TEX264-dependent processing of TOP1cc was hypothesised to occur through ATR-induced local disruptions to the interphase nuclear membrane, likely through phosphorylation and removal of lamin A/C. Interestingly, lamin A/C was detected in the lysosome under PARP trapping conditions in a leptomycin B-resistant manner (Fig. S6B). This fits the hypothesis that trapped PARP1 exits the nucleus through ATR-induced local disruptions in lamin A/C architecture.

### Disruption of the TEX264-p97 autophagy axis causes PARPi-induced replication-associated DNA damage

Having established the role of p97-TEX264-mediated selective autophagy in removing PARPi-induced trapped PARP1, we next sought to explore how the disruption of this affects cells. Trapped PARP1 causes increased replication stress, DSBs^107–109^ and replication-associated ssDNA gaps^110–112^ suspected to be caused by collision of the replication fork with trapped PARP1 and PARP1 trapping on unligated Okazaki fragments. Analysis of RNA-seq data comparing WT and TEX264^-/-^ cells treated with talazoparib shows significant differential expression of 60 and 37 genes related to DDR in HeLa and CAL51 cells, respectively (Fig. S7A, Tables S5 and S6), suggesting that TEX264-deficient cells experience altered DDR under PARPi treatment. In accordance with this, we observed increased levels of phosphorylated (p) RPA and pCHK1 in TEX264^-/-^ cells in response to PARPi (Fig. S7B), suggesting an increase in ATR signalling associated with replication stress. There were also heightened levels of the DDR markers γH2AX, RPA and 53BP1, with 1.4-, 2.1- and 1.8-fold increases, respectively (Fig. S7C and D). The same effect was observed in CAL51 cells (Fig. S7E and F), but only in response to talazoparib and not the weak trapping inhibitor veliparib, supporting that this DNA damage arises in response to accumulated trapped PARP1 when TEX264 fails to clear it from chromatin.

As the role of TEX264 in repair of trapped PARP1 appears to depend upon its role both as a selective autophagy receptor and p97 co-factor, we next explored how the interruption of these functions affects cellular response to PARPi. In TEX264^-/-^ cells complemented with TEX264^WT^, talazoparib-induced RPA and γH2AX levels were restored to those observed in WT cells (Fig. S8A and B). However, mutation of either the p97-interacting SHP motif or the LC3-interacting LIR domain failed to restore RPA and γH2AX levels (Fig. S8A and B). We observed the same effect on DNA damage foci when we inhibited p97 with CB-5083 (Fig. S8C and D) or autophagy through ATG7 depletion (Fig. S8E-G). It is worth noting that we did not detect an increase in PARPi-induced γH2AX foci under p97i conditions, as CB-5083 is known to impair ATM kinase activity^113^, the central kinase for phosphorylation of H2AX (γ-H2AX) in response to DNA damage^114^. Altogether, this follows our earlier data showing increased sensitivity to talazoparib by chemical or genetic inhibition of autophagy (Fig. 1D and E) and previously published work showing p97i hypersensitises cells to PARPi^39^.

### Autophagy is essential to clear PARP1 aggregates induced by trapping

Cells exhibit increased replication stress and DNA damage when trapped PARP1 is not efficiently cleared by TEX264-mediated autophagy. Previous studies have shown that p97, through its UFD1 cofactor, removes trapped PARP1 from chromatin via a proteasome-dependent pathway (p97-UFD1-proteasome)^39^. This raises the question of why the newly identified p97-TEX264-autophagy pathway is also important for cell survival. Notably, disrupting the p97-UFD1 pathway, either by UFD1 depletion or CB-5083 treatment, further increases the sensitivity of TEX264-/- cells to talazoparib (Fig. 6A and B), supporting that the p97-UFD1-proteasome pathway runs in parallel to the p97-TEX264-autophagy-mediated pathway. Moreover, the additive effect of TEX264 inactivation in cells treated with p97 inhibitors suggests that TEX264 may also play a role in processing trapped PARP independently of p97 (Fig. 6B). This also highlights that the clearance of PARP1 from chromatin by p97 alone is insufficient to promote cell survival. A key function of autophagy is in the clearance of protein aggregates^115^, and misfolded p97 substrates are thought to accumulate as aggregates^116, 117^ if not properly processed. In fact, camptothecin, a TOP1 inhibitor, was previously shown to induce aggregated TOP1 at doses where TOP1cc clearance is dependent upon autophagy^49^. Using the proteostat dye, which stains aggregated proteins, we were able to confirm by both FACS and immunofluorescence that aggregates also accumulate upon sustained treatment with talazoparib (Fig. 6C, D, and S9C). Accumulation of aggregates is sustained when autophagy is inhibited with bafilomycin A1, and they are observed to co-localise with LAMP1 (Fig. 6D), supporting that PARPi-induced aggregates are degraded by autophagy. Purification of insoluble aggregates showed that they contain high levels of PARP1, with the level of aggregated PARP1 increased substantially when talazoparib treatment was combined with bafilomycin A1 (Fig. 6E). Importantly, the PARP1 aggregates were not formed when cells expressing PARP1 that can not bind DNA/chromatin (PARP1-KS)^(ref)^. Overall, this indicates that autophagosomal processing of trapped PARP1 is key to preventing the accumulation of cytotoxic aggregated PARP1, and these PARP1 aggregates depend on PARP1 binding/trapping to chromatin.

**Figure 6:**
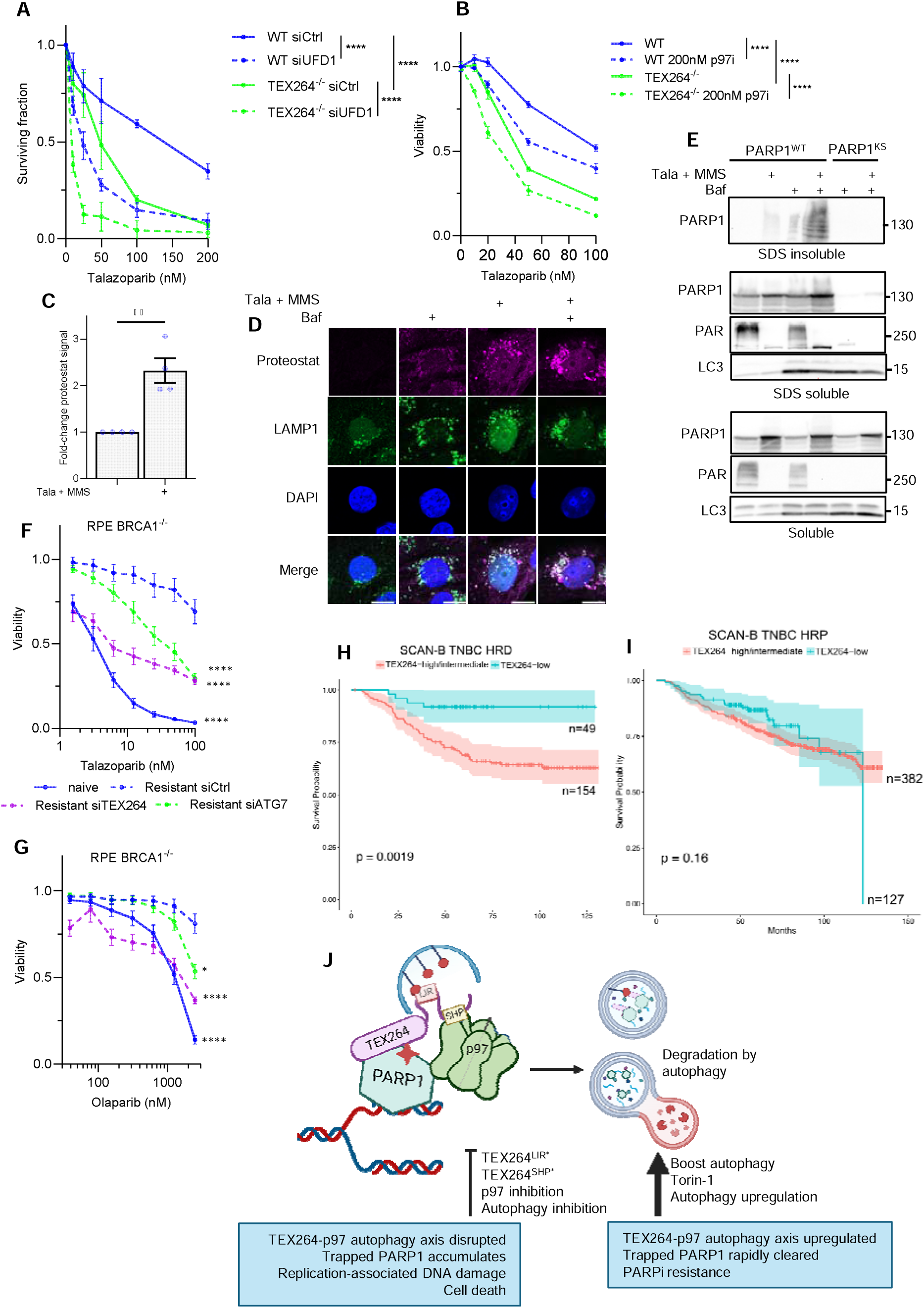
Loss of autophagy-dependent clearance of trapped PARP1 leads to accumulation of cytotoxic PARP1 aggregates and can overcome acquired PARPi resistance. **(A)** Colony formation assay in HeLa WT of TEX264^-/-^ cells depleted of UFD1 using siRNA and treated with talazoparib for 24 hrs. Statistical analysis by two-way ANOVA. **(B)** Cell viability by resazurin assay in HeLa WT or TEX264^-/-^ cells treated with talazoparib with or without p97 inhibitor CB-5083 (200nM) for 24 hrs, followed by 48 hrs recovery. Statistical analysis by two-way ANOVA. **(C)** Levels of protein aggregates measured by FACS using proteostat dye in cells treated with talazoparib and MMS for 2 hrs, then talazoparib alone for 18 hrs. Fold-change is compared to the untreated. Data is from 4 biological repeats with statistical analysis by Student’s unpaired t-test. **(D)** As in (C) but with aggregates visualised by immunofluorescence, showing accumulation of aggregates that co-localise with LAMP1 under trapping conditions when combined with bafilomycin A1 treatment. **(E)** Fractionation of CAL51 cells stably expressing PARP1^WT^ and PARP1^KS^ into the soluble fraction, SDS soluble fraction containing chromatin and SDS insoluble fraction containing protein aggregates. Immunoblotting determined PARP1 levels in each fraction. **(F and G)** Cell viability measured by resazurin assay in RPE TP53^-/-^ hTERT BRCA1^-/-^ cells, either naïve or resistant to olaparib. Resistant cells are depleted of TEX264, ATG7 or a luciferase control. Treatment is with **(F)** talazoparib or **(G) o**laparib for 6 days. 12 technical replicates across 4 biological repeats with statistical analysis by two-way ANOVA. **(H and I)** Kaplan-Meier plots of SCAN-B TNBC (n=712) in HRD and HR proficient tumours (n=203, 509 respectively). HR status was annotated using a 228 gene expression signature from Jacobson et al 2023 and compared against TNBC from TCGA-BRCA (n=312) with known HRD status. P-value was calculated using the log-rank test. **(J)** Model of the TEX264-p97-autophagy axis in trapped PARP1 repair. Inhibition of this process leads to cell death, whilst boosting this pathway results in cells developing resistance to PARPi.

### The TEX264-p97 autophagy axis is relevant to PARPi resistance

PARPi resistance is a major clinical concern. To explore if TEX264-mediated selective autophagy (nucleophagy) of trapped PARP1 may be relevant in the context of PARPi resistance, we used BRCA1^-/-^ cells that had acquired resistance to olaparib. For this, we used RPE1 TP53⁻ /⁻ hTERT BRCA1⁻/⁻ cells that had acquired resistance following prolonged exposure to olaparib. When compared to olaparib-naïve cells, we observed a striking resistance to both talazoparib (Fig. 6F) and olaparib (Fig. 6G), as expected. Interestingly, depletion of either TEX264 or ATG7 in resistant cells considerably re-sensitised them to both PARPi, with the strongest effect observed with talazoparib treatment, the more potent trapper of the two PARPi (Fig. 6F and G, S9A and B). This implies that impairing the clearance of trapped PARP1 by TEX264-mediated selective autophagy could partially overcome PARPi resistance. Given that the TEX264-nucleophagy axis is important for genome stability and cell survival in response to trapped PARP1, and BRCA1-deleted yet PARPi-resistant cells are hypersensitive to TEX264 inactivation, we asked whether this pathway has clinical relevance. We analysed RNA seq data from the SCAN-B^118^ cohort of 7743 breast cancer patients to examine the relationship between TEX624 expression and the homologous recombination status of these cancers. We focused on TNBC as previous reports have observed that trends in survival and homologous recombination status differ between TNBC and HER2+ and ER+ subtypes^118, 119^. Although there were no overall survival differences between homologous recombination proficient (HRP) and homologous recombination deficient (HRD) TNBC breast cancer patients (Fig. S10A), there was a strong and significant survival difference in HRD patients related to TEX264 expression (Fig. 6H,S10B-C), absent in HRP patients (Fig. 6I, S10D). HRD-breast cancer patients with low TEX264 mRNA expression showed about 28% better long-term survival, for example, after 10 years. These correlational, clinical data support our findings that a functional TEX264-nucleophagy pathway (e.g., high TEX264 expression) enables breast cancer cells to survive, and is associated with reduced overall patient survival (Fig. 6J).

## Discussion

Trapped PARP1 lesions induced by PARPi are highly cytotoxic and underlie the use of these drugs in treating HR-defective cancers. However, the rapid development of PARPi resistance poses a serious clinical challenge. Understanding the dynamics and regulation of trapped PARP1 repair is key to improving understanding of how resistance develops. Through three approaches, using new and published data from RNA-seq, mass spectrometry and CRISPR screens, we identified a direct role of selective autophagy in response to PARPi (Fig. 6J). Specifically, the SAR, TEX264, acts in tight coordination with the autophagy regulator LC3 and the ATPase p97 to directly process trapped PARP1 from chromatin and govern its delivery to the lysosome. This process depends on SUMOylation and ubiquitination, but is independent of the SUMO-dependent E3-ubiquitin ligase RNF4, which was previously shown to regulate trapped PARP1 degradation by the proteasome, indicating unknown E3 ligase(s) are involved. Defects in TEX264 interaction with LC3 or p97, by mutation of LIR or SHP domains, respectively, impair PARP1 delivery to the lysosome, cause increased PARPi-induced DNA damage and sensitise cells to talazoparib, showing it acts as a SAR and p97 co-factor in this pathway. Inhibition of this pathway causes sensitisation to PARPi through accumulation of cytotoxic aggregates of unfolded PARP1, and can re-sensitise resistant cells to both talazoparib and olaparib.

TEX264-driven selective nucleophagy of trapped PARP1 is tightly coupled to PARPi, with a firm reliance on PARPi and PARP1 trapping for localisation to the lysosome (Fig. 2B-D) and autophagy/TEX264 loss only causing sensitivity in a trapping-dependent context (Fig. 1D, 1G, 4A-F). Interestingly, other substrates of nucleophagy have been described under specific circumstances of oncogenic or genotoxic stress. Both lamin B1 and lamin A/C interact with LC3 and are degraded by autophagy in response to these stressors, inducing senescence^103, 104^. Similarly to trapped PARP1 (Fig. 2F-H), TOP2-DPCs and damaged DNA have been detected in nuclear buds that form upon etoposide treatment in an autophagy-dependent manner^120^. These examples demonstrate the relevance and selectivity autophagy of nuclear material, in responding to genotoxic stress, but do not implicate TEX264 as a SAR for this process or a direct processing of DNA lesions by autophagy. Our recently published work on TOP1cc^49^ was the first comprehensive example of autophagy clearing DNA damage, demonstrating that mammalian cells employ the p97-TEX264 system to remove TOP1cc to the lysosome. Therefore, we propose TEX264-mediated selective autophagy of DNA lesions (nucleophagy) is as a specialised DNA repair pathway for the removal of tightly bound chromatin protein lesions that are prone to form aggregates, TOP1cc^49^ and trapped PARP1 (Fig. 6). TEX264 was previously described to mediate selective autophagy of the ER (reticulophagy) through association with the ER membrane via its N-terminal leucine-rich region^51, 52^. In our case, where trapped PARP1 is the substrate of TEX264, TEX264 directly interacts with PARP1 through a motif located in its C-terminus (Fig. 5A-G). Due to the lack of knowledge surrounding the structure of TEX264, with no crystal structure available, the exact amino acids driving this interaction remain elusive.

The role of autophagy in cancer is complex, with different functions and effects depending on the type and stage of the disease^55^. There is equal confusion in the PARPi context, with conflicting roles of autophagy proposed^57^. Despite this, the evidence presented in our work suggests that autophagy is induced by PARPi treatment as a protective mechanism (Fig. 1), with most literature supporting this claim^56–63^. Until now, autophagy has been proposed to be induced by PARPi and act indirectly to promote cell survival, in line with other genotoxic agents^72^. PARPi-induced generation of reactive oxygen species (ROS) and upregulation of PTEN, a negative regulator of MTOR, enhanced autophagy, which acts to clear ROS, meaning its inhibition sensitised cells to olaparib through ROS accumulation. Further, it was shown that enhanced autophagy promotes HR, leading to increased BRCA1 and RAD51 recruitment to sites of PARPi-induced DNA lesions^87, 121^. Whilst we describe no alternative mechanism for how PARPi induce autophagy, we observe a direct role of cytoprotective nucleophagy in processing trapped PARP1, as no effect is observed in non-trapping conditions, with veliparib (Fig. 1F, 2D) or trapping-defective mutant PARP1^del.p.119^^K120S^ (Fig. S3A and B). The nucleophagy described here is likely activated in response to PARP1-bound DNA-induced aggregates stimulated by PARPi and the resulting trapping of PARP1 on chromatin. Our hypothesis is further supported by the observation that a PARP1 mutant unable to bind chromatin does not form aggregates (Fig. 6E). Of all PARPi, the most promising evidence for cytoprotective autophagy is associated with talazoparib^57^, the most potent trapper amongst clinically approved PARPi^14, 16^. Together, this strongly supports the discovery of a novel and direct role of nucleophagy in processing trapped PARP1.

Pre-clinical evidence in cell lines and patient-derived organoids of various cancer types shows combined treatment of PARPi and autophagy inhibitors as a promising therapeutic strategy^57^. Interestingly, after chronic treatment to induce olaparib resistance, breast cancer cell lines show upregulation of autophagy compared to their olaparib-naïve counterparts, with autophagy inhibition able to re-sensitise resistant cells to olaparib^63^, implying a potentially active role of autophagy in resistance. Autophagy upregulation has also recently been linked to olaparib resistance in pancreatic cancer cells, both *in vitro* and *in vivo*^122^. Whilst attractive, targeting autophagy in cancer therapy can be complex due to its pleiotropic and conflicting roles throughout tumorigenesis, metastasis and in the tumour microenvironment^123, 124^, as shown by mixed patient responses observed in clinical trials using chloroquine derivatives in combination with various therapies^123^. In our study, resistant cells generated by chronic olaparib treatment were re-sensitised to PARPi by knockdown of TEX264 or ATG7 (Fig. 6F and G), indicating that specifically inhibiting nucleophagy of trapped PARP1 could be used to overcome resistance whilst limiting off-target effects of broader autophagy inhibitors. Importantly, analysis of clinically available data strongly suggests that TEX264 expression levels play a significant role in the overall survival of HR-deficient breast cancer patients (Fig. 6H and I). These findings further emphasise the significance of the TEX264-nucleophagy pathway in our understanding of genome stability and highlight the potential use in predicting therapeutic responses in cancer patients. In conclusion, we identified p97-mediated TEX264-orchestrated nucleophagy as a specialised pathway for processing trapped PARP1 and maintaining genome stability in response to PARPi. Importantly, we have demonstrated that nucleophagy is directly involved in the clearance of PARPi-induced trapped PARP1, highlighting a potential mechanism of PARPi resistance that could be further explored in the clinic.

## Materials and methods

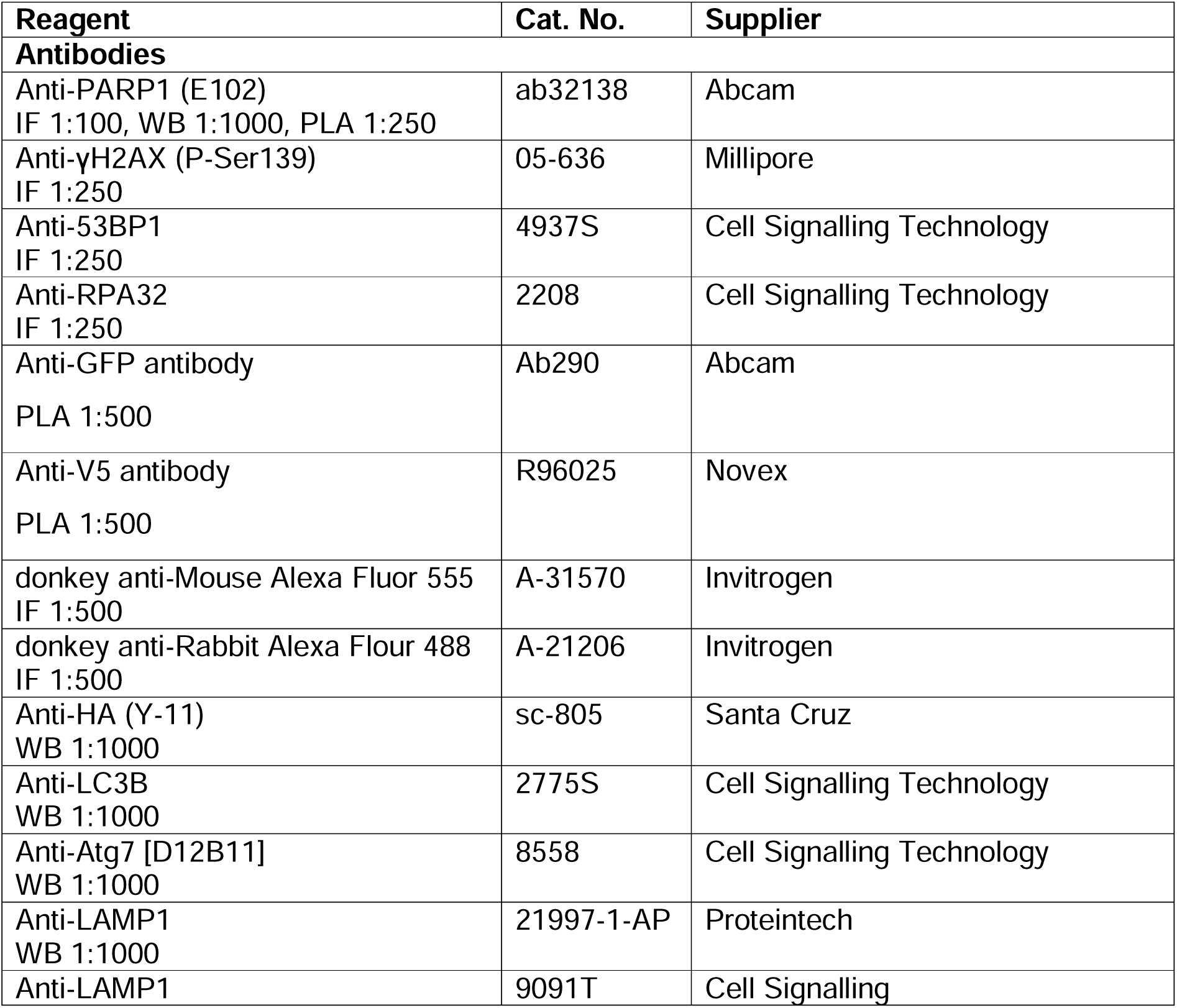
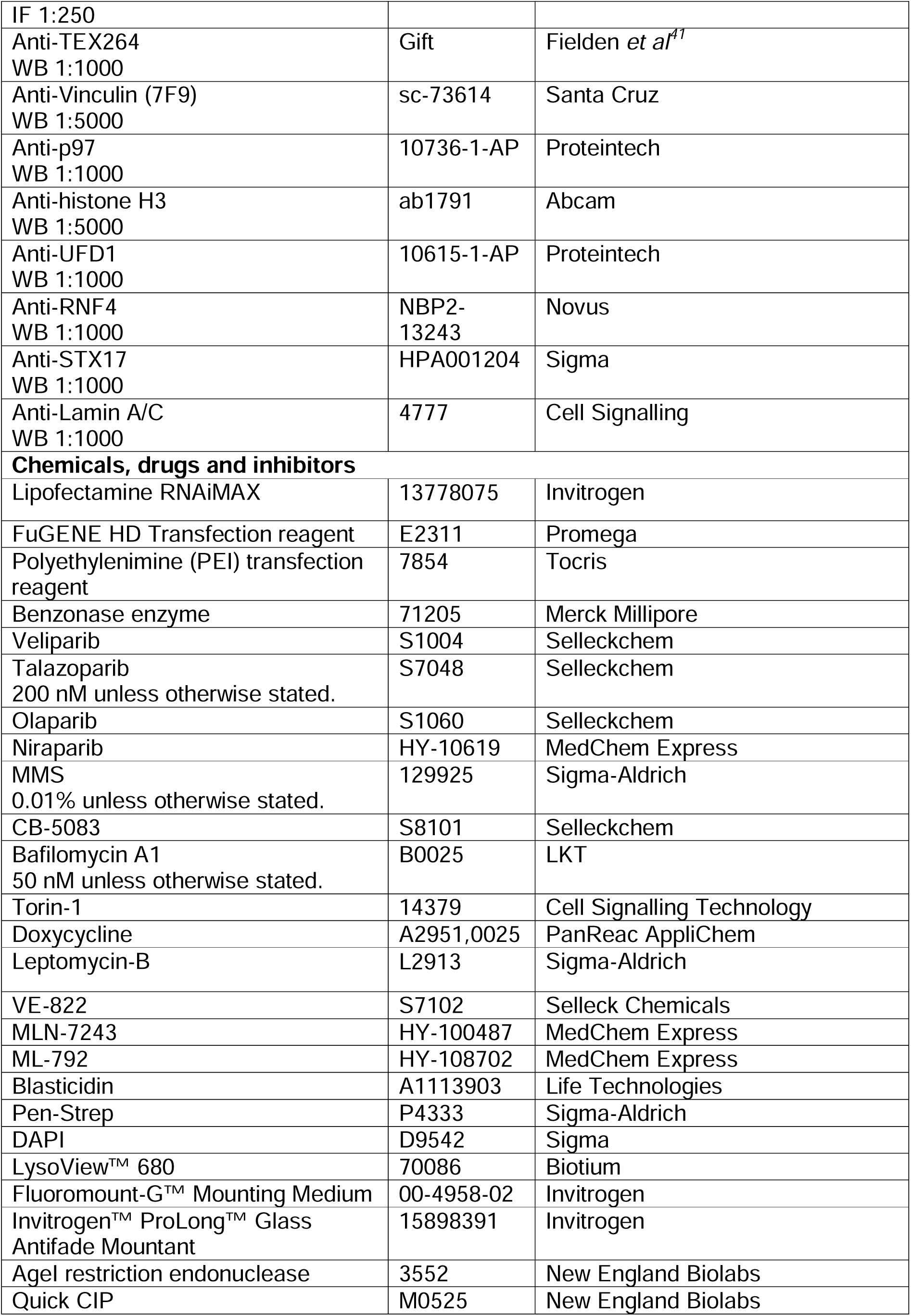
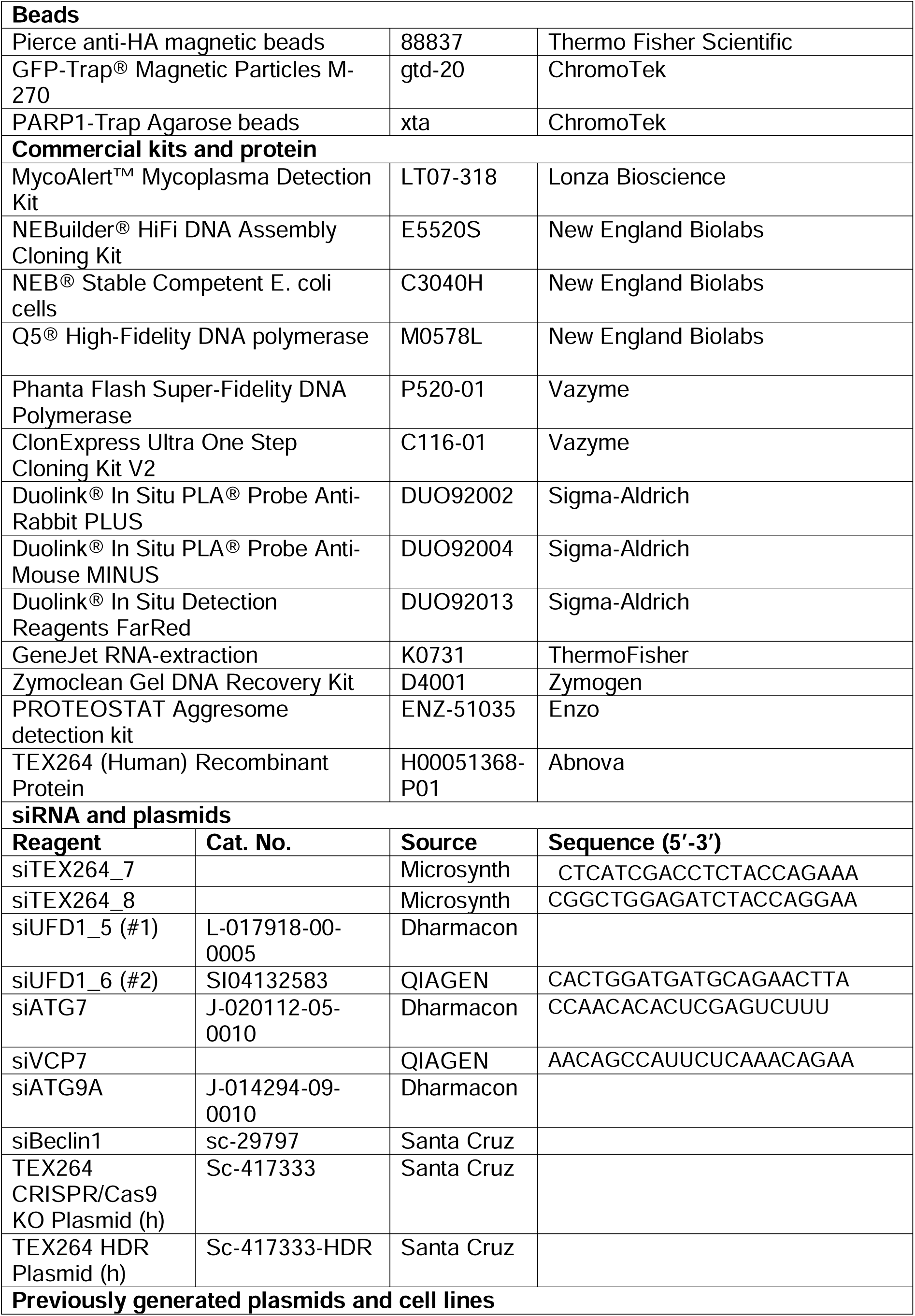
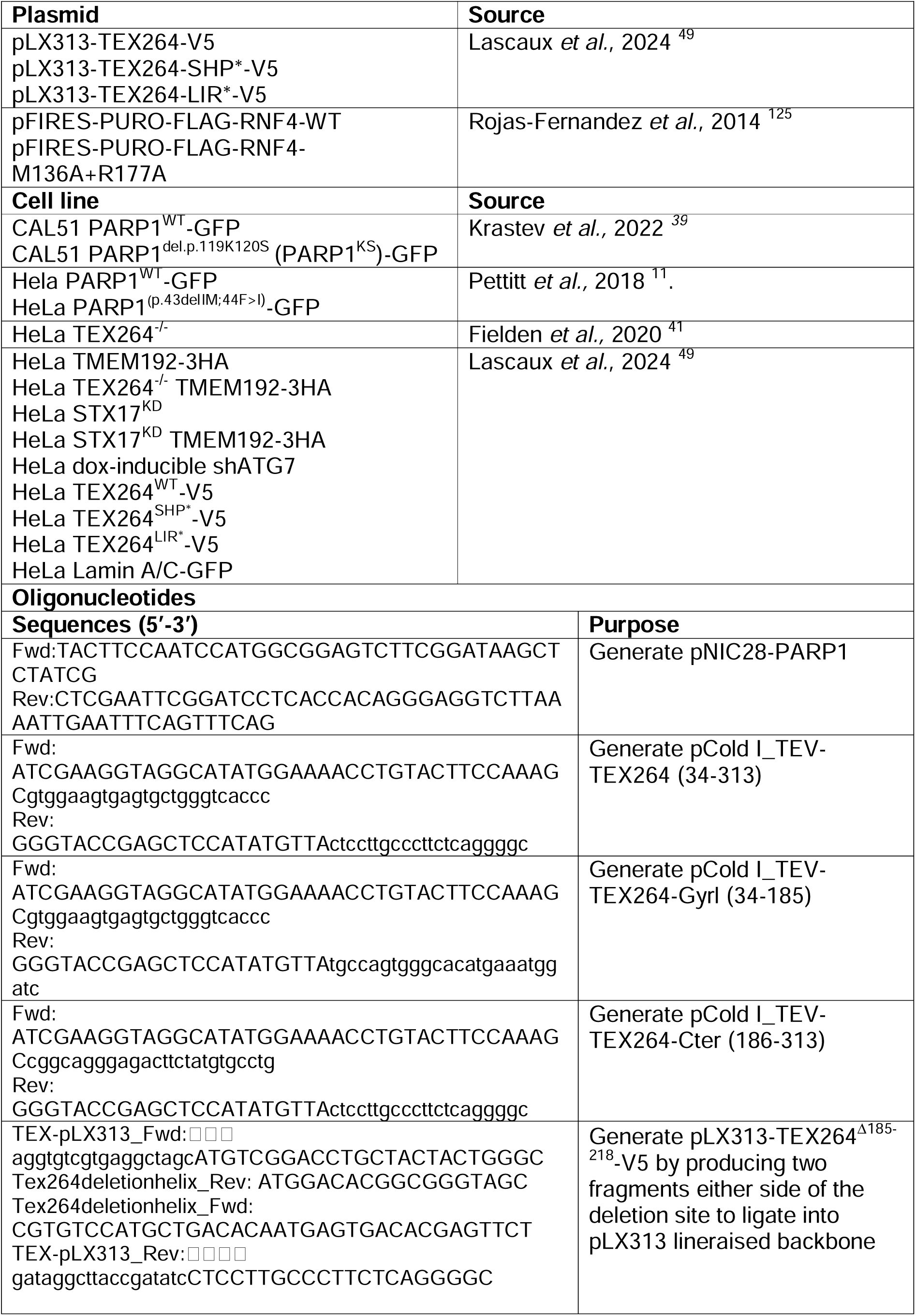
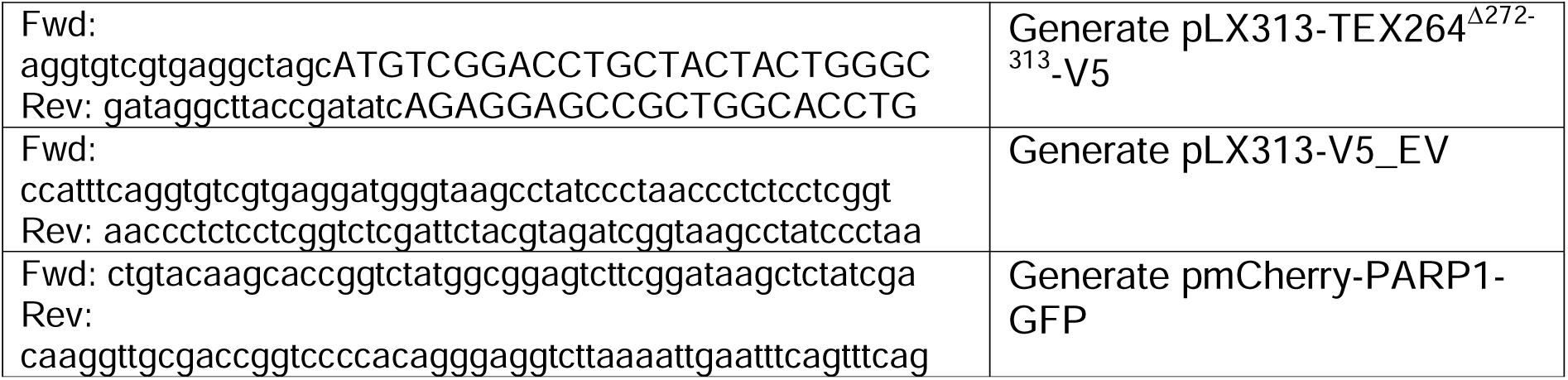

### Cell culture, transfection and drug treatment

CAL51 (DSMZ, ACC 302), HeLa (ATCC, CCL-2), MDA-MB231 (ATCC, Htb-26) and RPE TP53^-/-^ hTERT BRCA1^-/-^ were maintained in Dulbecco’s Modified Eagle Medium (DMEM), supplemented with 10% fetal bovine serum and 1x penicillin–streptomycin (Sigma-Aldrich). Cells were tested for mycoplasma regularly. Cells were transfected with siRNA using Lipofectamine RNAiMAX according to the manufacturer’s instructions, with experiments carried out 72 hours after transfection. Transient transfection with plasmids was carried out using FuGENE HD Transfection reagent for microscopy-based experiments and Polyethylenimine (PEI) transfection reagent for all other experiments, with both used according to the manufacturer’s instructions. Drug treatments were as indicated in the figure legends, with a vehicle control used where appropriate. Talazoparib was used at 200 nM, MMS at 0.01% and bafilomycin A1 at 50 nM unless otherwise stated.

### Generation of plasmids and stable cell lines

To generate pLX313-V5_EV, two overlapping oligos were ligated into a pLX313 backbone cleaved at the EcoRV/NHeI restriction digest sites to insert a STOP codon after the V5 tag. Ligation was performed using NEBuilder® HiFi DNA Assembly, then transformed into NEB® Stable Competent E. coli cells. Successfully transformed colonies were identified by colony PCR and sequenced by Source BioScience, Oxford, UK. pLX313-TEX264-V5 variant plasmids were generated in a similar way, using the oligos listed in the materials table. TEX264 fragments were PCR amplified using Q5® High-Fidelity DNA polymerase. Full-length PARP1 was cloned into NcoI and BamHI restriction sites in the pNIC28 vector in a similar way. TEX264 (34-313; 34-185; 186-313) carrying a TEV cleavage sequence on the 5’-end was cloned by the Phanta Flash Super-Fidelity DNA Polymerase. The fragment was then inserted into the pCold-I vector at the NdeI site using the ClonExpress Ultra One Step Cloning Kit V2. pmCherry-PARP1-eGFP was generated by the Genome Engineering & Transgenics facility at the MRC Wetherall Institute of Molecular Medicine. PARP1 cDNA was amplified from an existing plasmid and cloned by InFusion (Takara) into pmCherry-eGFP (Addgene, #86639) at AgeI restriction sites. InFusion reactions were transformed into Stable2 competent E.coli, and correct colonies were identified by AgeI digestion and single-molecule sequencing (Oxford Nanopore).

CAL51 TEX264^-/-^ were created by CRISPR/Cas9 knockout of TEX264 using TEX264 CRISPR/Cas9 KO Plasmid (h) and TEX264 HDR Plasmid (h). Plasmids were transfected into CAL51 cells using Fugene, and then the transfected cells were selected with 2 µg/mL puromycin. Single-cell colonies were isolated by limiting dilution and validated for TEX264 loss by Western Blot. Cells were induced to stably express TMEM192-3HA, TEX264^WT^-V5, TEX264^SHP*^-V5, TEX264^LIR*^-V5 or empty vector (EV)-V5 using lentiviral transduction with the following plasmids: pLJC5-Tmem192-3xHA (Addgene, 02930), pLX313-TEX264-V5, pLX313-TEX264-SHP*- V5, pLX313-TEX264-LIR*-V5, pLX313-V5_EV-V5. Lentiviral particles were generated in HEK293T cells by transfection with transfer plasmid for gene of interest, pAmphoR envelope plasmid and Δ8.2R packaging plasmid, which were both a generous gift from Vincenzo D’Angiolella, using PEI transfection. After 72 hours, viral particles were harvested, filtered and added to CAL51 PARP1^WT^-GFP, CAL51 PARP1^KS^-GFP, CAL51 TEX264^-/-^, HeLa TEX264^-/-^ cells, as required, with 16 µg/mL polybrene. Puromycin or hygromycin was used to isolate TEX264 or TMEM192-3HA transduced cells, respectively, before isolating and expanding single-cell colonies by limiting dilution. Colonies were screened by Western Blot and immunofluorescence to select cell lines that were stably expressing TEX264 variants at a similar level to the endogenous.

Human retinal pigment epithelial RPE1 BRCA1^−/−^ cells transduced with hTERT and TP53-deleted^126^ were cultivated in monolayers in DMEM supplemented with 10% foetal bovine serum in the presence of 2 μg/mL blasticidin. To generate PARP inhibitor-resistant cells, BRCA1^−/−^ RPE1 cells were grown in the presence of increasing doses of olaparib for 3 months. Briefly, cells were seeded to 50% confluency and initially treated with 20 nM olaparib. Cells were allowed to grow to 80-90% confluency, whilst fresh media supplemented with the drug was replaced every 2-3 days. Olaparib concentration was increased by 25% each time the cells were passaged to a final concentration of 76.25 nM olaparib. Viability assays were conducted to confirm resistance to PARP inhibitors.

The sources of previously generated plasmids and cell lines are listed in the materials table.

### Immunofluorescence

Immunofluorescence was carried out as previously described^39^ with detergent pre-extraction. Cells were seeded and grown on glass coverslips to 70-90% confluency. After 1 wash with PBS, pre-extraction buffer (25 mM HEPES, pH 7.5, 50 mM NaCl, 1 mM EDTA, 3 mM MgCl2, 300 mM sucrose and 0.5% (v/v) Triton X-100) was added on ice for 2-2.5 minutes. Cells were washed once with the same buffer without Triton X-100 before fixing for 15 minutes on ice with 4% formaldehyde in PBS. Coverslips were blocked with 5% BSA for 1 hr at 37°C, then sequentially incubated with antibodies diluted in 2.5% BSA for 1 hr at room temperature. Antibodies used were: anti-PARP1, anti-γH2AX, anti-RPA, anti-53BP1, donkey anti-Mouse Alexa Fluor 555, and donkey anti-Rabbit Alexa Fluor 488. Images were acquired using either the Andor Dragonfly confocal or Nikon Ni-E widefield microscopes and analysed with custom Cell Profiler pipelines.

### Western Blot

Standard protocols were used for SDS-PAGE and subsequent immunoblotting using either 0.22μm pore size PVDF (BioRad) or nitrocellulose (GE Healthcare) membranes to transfer proteins from homemade polyacrylamide gels.

### Cell survival assays

For colony formation assays, cells were seeded in 6-well plates at 1000 cells/well for WT and 1500 cells/well for TEX264^-/-^ cells. After 16 hours, cells were treated for 24 hours, then allowed to grow in recovery media for 6-10 days until colonies were 20-50 cells in diameter. Wells are washed with PBS, then fixed in 100% methanol for 10 minutes before staining in crystal violet (1.23 mM crystal violet, 1% formaldehyde, 1% methanol, 1x PBS). Colonies were scanned and counted using GelCount (Oxford Optronix). For resazurin assays, 500-1000 cells/well were seeded in black well flat-bottom 96-well plates. The following day, treatment was added for the time described in the figure legends. After treatment was complete, the media was replaced with fresh media containing 30 µg/mL resazurin for 4-6 hrs. Resazurin media was added to 3 empty wells to serve as a blank control. Fluorescence was measured at 570 nm using a plate reader. The average fluorescence detected in the blank sample was subtracted from each reading to normalise for background signal. For both assays, all conditions were in technical triplicate. Resazurin assays were more commonly used when experiments required depletion by RNAi, as this often affected colony formation. The shorter time course of resazurin assays was also more suited to depletion-based experiments, as low protein levels could be maintained more reliably than in longer colony formation assays.

### Chromatin fractionation and co-immunoprecipitation

Chromatin fractionation and co-immunoprecipitation were performed as previously described^39^. Briefly, sub-confluent cells were harvested in PBS containing 3mM EDTA. Nuclear pellet was isolated by lysis in buffer A (10 mM HEPES pH 7.45, 10 mM KCl, 340 mM sucrose, 3 mM EDTA, 10% glycerol, protease and phosphatase inhibitors, NEM, 0.1% triton X-100), then chromatin was isolated in buffer B (3 mM EDTA, 0.2 mM EGTA, 5 mM HEPES pH 7.9, protease and phosphatase inhibitors, NEM). Soluble chromatin was recovered by digestion in benzonase buffer (50 mM Tris-HCl pH 7.9, 100 mM NaCl, 10 mM MgCl_2_, 125 U/mL benzonase). An input sample was taken, and 50µg/ml ethidium bromide was added to the remainder. PARP1-GFP was captured from the soluble chromatin fraction on GFP-trap beads, previously blocked in 5% BSA. Beads were washed 3 times for 15 minutes with IP wash buffer (50 mM Tris-HCl pH 7.4, 150 mM NaCl, 0.5 mM EDTA, 0.05% Triton X-100, protease and phosphatase inhibitors, NEM), before elution with Laemmli buffer.

### Biochemical isolation of aggregates

The isolation of aggregates was performed as previously described ^49^. Briefly, CAL51 cells were treated for 1 hr with talazoparib and MMS, followed by talazoparib with or without bafilomycin A1 for 18 hrs. Cells were harvested and lysed with benzonase to collect the soluble fraction. The pellet was washed with a buffer containing 1.5% SDS to collect the SDS-soluble fraction. The SDS insoluble fraction was solubilised with 100% formic acid and sonication, before evaporating the formic acid and resuspending the pellet in Laemmli buffer for running by Western blot.

### Proximity ligation assay

Proximity ligation assay was performed using the Duolink® In Situ PLA® kits, following the manufacturer’s protocol. CAL51 cells stably expressing both PARP1-GFP and TEX264-V5 were seeded and grown on glass coverslips to 70-90% confluency. After treatment, cells were fixed with 4% formaldehyde in PBS for 10 minutes, then permeabilised with 0.25% Triton X-100 for 10 minutes. After 3 washes in buffer A (150 mM NaCl, 10 mM Tris pH 7.4, 0.05% Tween 20), cells were blocked in the provided blocking reagent. Further washes in buffer A were followed by incubation with anti-GFP and anti-V5 primary antibodies (1:500). Coverslips were then incubated with PLA PLUS and MINUS probes. Cells were incubated with ligase followed by polymerase, with washes in buffer A between each step. Coverslips were then washed twice with wash buffer B (100 mM NaCl, 250 mM Tris pH 7.5) before staining with DAPI (1:1000) for 10 minutes. Finally, coverslips were washed twice with buffer A and once with 0.01x buffer B before mounting using ProLong™ Glass Antifade Mountant. Images were acquired using a TCS SP8 laser scanning confocal microscope (Leica, Germany). A 63x 1.2NA water immersion objective lens was used for acquisition, with the confocal pinhole size set to 111.4um. Images were scanned at 2048*2048, with a pixel size of 90nm. Analysis was carried out using ImageJ and a bespoke pipeline on CellProfiler.

### Immunoprecipitation of intact lysosomes (LysoIP)

Lyso-IP was performed as previously described^88^. Briefly, cells expressing TMEM192- were harvested and washed in KPBS (136 mM KCl, 10 mM KH_2_PO_4_, adjusted pH 7.25 with KOH) before an input sample was taken. Cells were homogenised with 15 strokes in a Dounce homogeniser, then centrifuged at 1000g for 2 minutes. Supernatant containing organelles, including lysosomes, was loaded onto anti-HA magnetic beads and incubated for 15 minutes on a rotating wheel at 4°C. Beads were washed 5 times in KPBS, then eluted in Laemmli for analysis by Western blotting. Quantification of band intensity was performed using ImageJ. PARP1 levels in the lysoIP fraction were normalised by dividing by the HA level in the lysoIP fraction. Each set of biological repeats was then divided by one condition to display the PARP1 level as a fold-change.

### mCherry-PARP1-GFP assay

For fixed imaging, cells transfected with pmCherry-PARP1-GFP were seeded on glass coverslips, as for immunofluorescence. After treatment, coverslips were washed once in PBS and fixed with 4% formaldehyde in PBS for 15 minutes, then washed 3 times with 0.01% BSA in PBS, before incubation with DAPI for 30 minutes in the dark. After a further 3 washes, coverslips were mounted on glass slides using Invitrogen™ ProLong™ Glass Antifade Mountant. Images were acquired using a TCS SP8 laser scanning confocal microscope (Leica, Germany), as described for proximity ligation assays, and analysed using ImageJ. Quantification was performed manually by observing the number of red cytosolic puncta per cell, with only puncta in proximity to the nucleus counted.

For live cell imaging, cells transfected to express mCherry-PARP1-GFP were seeded in 35 mm glass-bottom dishes. LysoView™ 680 was added 20 minutes before beginning the live imaging assay, FluoroBrite™ DMEM (A1896702, Gibco) to reduce background fluorescence. Confocal images were captured on an Olympus IXplore Spin-SR microscope using a 50um pinhole spinning disc. A 60x/1.3NA Lens was used, and images were obtained using a Hamamatsu ORCA fusion BT camera. 12um Z-stacks were captured using a z spacing of 0.8um, over a 4-hour time course at 35-second intervals. Rendering was performed using the TrackMate plugin on ImageJ.

### Proteostat aggregates assay

For FACS, cells in 6-well plates were treated with talazoparib and MMS for 2 hrs, followed by treatment with talazoparib only for 18 hours. After harvesting with trypsin, cells were washed in PBS, then the cell pellet was re-suspended in 200 µL PBS. This was added dropwise into 1 mL of 4% formaldehyde with slow vortexing. After 30 mins fixation at room temperature and one wash with PBS, cells were resuspended and added as for fixation into 1 mL of permeabilisation buffer (0.5% Triton X-100, 3 mM EDTA, pH 8.0) in PBS. After 30 minutes on ice, cells were once with PBS. Cells were transferred through the cell strainer cap of a FACS tube (352235, Corning) to remove debris, then centrifuged at 800 x g for 10 minutes. The cell pellet was resuspended in 500 µL of PROTEOSTAT® Aggresome Red Detection Reagent diluted 2,500-fold in 1x Assay Buffer and incubated for 30 minutes in the dark. Samples were analysed in the FL3 channel of an LSR Fortessa.

Immunofluorescence of aggregates was carried out in a very similar way to standard immunofluorescence, with the proteostat dye applied to coverslips after fixation and permeabilisation.

### *In vitro* immunoprecipitation

The purification of required proteins is described in detail in the supplementary note. PARP1-trap beads were washed in permissive buffer (50 mM HEPES, pH 7.4, 500 mM NaCl, 0.01% Triton X-100, 1 mM TCEP-HCl), followed by blocking in the same buffer containing 5% BSA for 1 hr. After two further washes in permissive buffer, PARP1 was loaded in 150 µL permissive buffer to a concentration of 2.5 µM and incubated with rotation for 1 hr at 4°C. Beads were washed 3 times with permissive buffer, followed by one wash with a more stringent buffer containing 0.1% Triton X-100. TEX264 was incubated with blocked beads in parallel to remove proteins prone to non-specific binding. TEX264 in the supernatant was added to PARP1-bound beads in 150 µL permissive buffer to a concentration of 2.5 µM, and an input sample was collected before incubating with rotation for 2 hrs at 4°C. Beads were washed twice with permissive buffer, then 3 times with stringent buffer before resuspending in Laemmli buffer containing 100 mM DTT for analysis by Western blot.

### Hydrogen-Deuterium eXchange mass spectrometry

#### Sample Preparation

We designed Hydrogen Deuterium eXchange Mass Spectrometry (HDX-MS) experiments to map the PARP1 interaction interface on TEX264, whilst also obtaining structural and mechanistic insights into the complex formation itself. TEX264 protein stocks were provided at 20 μM in 50 mM TRIS, 10 mM reduced glutathione. PARP1 protein stocks were provided at 19 μM in 50 mM HEPES, 150 mM NaCl, 0.5 mM TCEP. TEX264 was incubated with (holo-state) and without (apo-state) PARP1 for 60 min at room temperature to enable the complete formation of the complex. A molar ratio of 3:1 (PARP1:TEX264) was used. Samples were diluted over the course of the labelling experiment to achieve a final TEX264 concentration of 20 pmol on column.

#### Data Acquisition

We implemented an HDX-MS strategy similar to that which we recently described^127^. Briefly, we prepared a deuterium oxide D2O (99+ %D, Cambridge Isotope Laboratories, Tewksbury, MA) labelling buffer supplemented with identical buffer conditions to those of the protein stocks. The pH was corrected to pD 7.42 (pD=pH+0.4). A quenching buffer of 0.8% formic acid, pH 1.08 in H2O, was prepared. For the labelling reaction, samples were diluted in the deuterium labelling buffer in a 1:10 ratio to achieve a final excess D2O concentration of 90%. Labelling time points of 0.5, 10, and 60 min were sampled at 20°C, with matching non-deuterated controls in H2O buffer. A minimum of triplicate analyses was obtained for each time point and condition. At the end of each labelling time, the reaction was stopped by adding quench buffer (1:1 ratio) to reach a final pH of 2.49. Protein samples were digested with a pepsin and protease XIII acidic dual protease column (2.1 x 3.0 mm; NovaBioAssays, MA) at 8°C for 3 min. Peptides were subsequently trapped on a 1.0 mm x 5.0 mm, 5.0 µm trap cartridge (Thermo Scientific™ Acclaim PepMap100) for desalting using a flow rate of 150 µL/min. Peptides were separated on a Thermo Scientific™ Hypersil Gold™ column (50 x 1 mm, 1.9 μm, C18) by a linear gradient of 5% to 40% Buffer B (A: water, 0.1% FA; B: ACN, 0.1% FA) and a flow rate of 40 µL/min. To limit peptide carry-over, a protease wash of 2 M guanidine, 0.8% formic acid, pH 2.3 in H2O was performed after each injection. To minimise back-exchange, the LC system was maintained at a temperature of 1.5°C. Labelling, quenching, and online digestion steps were performed with the aid of an automated HDX robot from Trajan Scientific and Medical, which was guided by Chronos software (version 5.4.1). Samples were acquired in MS1 mode on a Thermo Scientific™ Orbitrap Exploris™ 480 Hybrid™ mass spectrometer.

#### Data Analysis

In the first instance, an unspecific digested database of non-deuterated TEX264 and PARP1 peptides was generated in BioPharma Finder (version 5.2) using a data-dependent and targeted HCD-MS2 acquisition regime. Processing and curation of the labelling data were performed with the aid of HDExaminer version 3.4.2 (Trajan Scientific and Medical). The charge state with the highest quality spectra for all replicates for each peptide across all HDX-MS labelling times was used in the final analysis. Alpha-fold were used to compare apo- and holo-states of TEX264 and the impact of PARP1 binding. Significant differences observed at each residue of the protein were used to map HDX-MS consensus effects (based on overlapping peptides) onto the AlphaFold model of TEX264. All MS raw files were deposited in ProteomeXchange (https://www.proteomexchange.org/) under the unique identifier PXD071389. **Token:** hyRVOEKeaKEv. Alternatively, the data can also be accessed by logging in to the PRIDE website using the following account details: Username. Password: WLt6XQ4zTtcd

#### RNA extraction and sequencing

For RNA-sequencing experiments, RNA was extracted from HeLa and CAL51 WT and TEX264^-/-^ using the GeneJet RNA purification kit, carried out according to the manufacturer’s instructions. RNA was quantified by nano-drop to ensure the concentration was higher than 20 ng/µL, and both the 260/230 and 260/280 ratios are above 2.0 to indicate sufficient purity. 3 biological repeats were sent to Novogene for sequencing, RNA sample quality control, mRNA library preparation (polyA enrichment), Illumina Sequencing and bioinformatic analysis. Bioinformatic analysis performed by Novogene included: (i) data quality control and filtering, (ii) mapping to reference genome GRCh38/hg38, (iii) gene expression quantification and correlation analysis, (iv) differential expression analysis, (v) enrichment analysis and (vi) gene set enrichment analysis.

#### Genome-wide CRISPR/Cas9 screens analysis

Genome-wide CRISPR/Cas9 screens were previously published^31, 76^. For the analysis in this study, quality control was performed using R software (R Core Team, 2024). Sequence alignment and enrichment analysis (day 0 vs PARPi-treated population) were carried out using the MAGeCK Maximum Likelihood Estimation (MLE) module^128^ and the R package MAGeCKFlute^129^. The dataset of MAGeCK MLE analysis results of the CRISPR/Cas9 screen on RPE1-h*TERT* cells was extracted from the Supplementary Table 1 of Noordermeer *et al.,* (2018)^31^. Datasets were robustly z-normalised and filtered against an untreated control population. Genes were considered depletion hits only when scoring at least two standard deviations under the median of each screen, under the treated condition and not in an untreated control population. Functional Enrichment analysis was performed using the R package *clusterProfiler*^130^ on the following databases: KEGG Pathways^131^, Reactome^132^, Gene Ontology: Biological Process^133^ and Complex^134^.

#### Survival analysis based on TEX264 expression and HRD status

The unadjusted RNAseq gene counts from the SCAN-B cohort^118^ were obtained (https://data.mendeley.com/datasets/yzxtxn4nmd/4), and read count normalisation using DESeq2 (v1.38.3) was done. The analysis was limited to TNBC (n = 712) in the SCAN-B cohort, as HRD survival trends were previously observed to differ between TNBC and HER2+ and ER+ subtypes^118, 119^.

To annotate the HRD status of each patient in the SCAN-B cohort, a 228 HRD gene set from Jacobson 2023^135^ was used, to carry out non-negative matrix factorisation (NMF package v0.26, method = brunet, nrun = 100) to identify gene expression signatures associated with HRD. In addition, FPKM-normalised gene expression data from TNBC patients (n = 312) in the TCGA-BRCA cohort were obtained from the GDC Data Portal (https://portal.gdc.cancer.gov/) to correlate HRD gene expression signatures in tumours previously annotated^135^ as HRD and HRP. Signature tnbc.2 correlated with HRD tumours in the TCGA cohort (Suppl Fig. 10E) and matched signature scanb 4 with a Pearson correlation of 0.86 (Suppl Fig. 10F) in the SCAN-B cohort. SCAN-B TNBCs with proportion of scanb.4 signature weights > 0.25 were annotated as HRD.

SCAN-B tumours were stratified using *TEX264* expression based on quartiles, with the highest quartile as TEX264-high, the lowest quartile as TEX264-low and the remaining tumours labelled as TEX264-intermediate.

Survival analysis was performed in R using the survival package (v3.5-5) with overall survival (OS) as the endpoint. Survival curves were compared using Kaplan-Meier curves generated using the survminer (v 0.5.0) package, and statistical tests were performed using the log-rank test.

### Data representation and statistical Analysis

Image data analysis and representation were carried out using CellProfiler™ (Broad Institute, https://cellprofiler.org/) and ImageJ (NIH, https://imagej.net/Fiji/Downloads). Images are shown with scale bars of 10 µm unless otherwise stated. Graphs were plotted, and statistical analysis was performed using Prism v10 (GraphPad Software, https://www.graphpad.com). All experiments were performed at least 2 times, with the number of replicates indicated in the figure legends. Error bars show SEM. Tukey box plots are standard, where the centre line equals he median, the box indicates the interquartile range, and whiskers indicate 1.5x interquartile range. Statistical tests (Student’s t-test, one-way ANOVA, two-way ANOVA) used are indicated in the figure legends. Asterisks are used to indicate p values (p>0.05 = ns, p≤0.05 = *, p≤0.01 = **, p≤0.001 = ***, p≤0.0001 = ****).

## Data Availability

All data generated, analysed and used in this study are included in this published article and its supplementary Information. Source data from published CRISPR screens is available in the European Nucleotide Archive under accession number PRJEB74933 (https://www.ebi.ac.uk/ena/browser/view/PRJEB74933) for data from Dibitetto *et al.,* (2024)^76^ and in Supplementary Table 1 of Noordermeer *et al.,* (2018)^31^. Published mass spectrometry data^39^ is available in the ProteomeXchange Consortium via the PRIDE partner repository (dataset identifier PXD024337). Source data for RNA-seq experiments is deposited in Gene Expression Omnibus and can be accessed using the GEO accession number GSE277366. To review GEO accession GSE277366: Go to https://www.ncbi.nlm.nih.gov/geo/query/acc.cgi?acc=GSE277366

Enter token ctmpawmobduxpqh into the box. All plasmids generated in this manuscript will be deposited in Addgene (https://www.addgene.org/browse/). Please request these plasmids directly from Addgene. Cell lines created in this study are available from the corresponding author upon reasonable request.

## Author contributions

Conceptualisation, K.R. and G.H.; Methodology, Formal Analysis and Investigation, G.H.; S,T., P.L., S.K., I.T., W.S., C.H., J.L., D.O., A.P., A.W.T.N., Y.L., and K.R. Validation, G.H., P.L., C.H., J.L. and K.R.; Resources, G.H., P.L., M.G.F., G.G., R.D., N.R., I.M., R.F., S.R., D.B.K., C.J. L. and M.T.; Data Curation, all authors; Writing – Original Draft, G.H.; Writing – Review & Editing, G.H. and K.R.; Visualisation, G.H.; Supervision, K.R.; Funding Acquisition, G.H., P.L., and K.R.

## Acknowledgements

The authors would like to thank all members of the Ramadan Laboratory, Oxford, for their help in data interpretation and critical comments during the execution of this study. We would like to acknowledge the Genome Engineering & Transgenics facility at the MRC Weatherall Institute of Molecular Medicine for their assistance in molecular cloning of plasmids. This work received support from the Medical Research Council programme (Grant Ref. MR/X006409/1), Breast Cancer Now (Grant Ref. 2022.11PR1570), Ministry of Education-Start-Up Grant, Singapore, Toh Kian Chui Distinguished Professorship Award /LKC Medicine, Singapore to K.R., the Luxembourg National Fund (Grant Ref. 14548187) studentship awarded to P.L., Medical Research Council studentship awarded to G.H, Four Diamonds Pediatric Cancer Research Funds to N.R., grant PID2022-139691OB-I00 funded by MCIN/AEI/10.13039/501100011033 and ERDF A way of making Europe (European Union) to R.F., Wellcome Trust (224361/Z/21/Z) and John Black Foundation to I.M. Work in M.T. laboratory was supported by Cancer Research UK Program Award (DRCPGM\100001 (M.T.). Financial support for the S.R. laboratory came from the Swiss National Science Foundation (320030M_219453) and the Office of the Assistant Secretary of Defense for Health Affairs through the Ovarian Cancer Research Program under Award No. W81XWH-22-1-0557.

## Declaration of interest

C.J.L. makes the following disclosures: receives and/or has received research funding from: AstraZeneca, Merck KGaA, Artios, Neophore, FoRx. Received consultancy, SAB membership or honoraria payments from: FoRx, Syncona, Sun Pharma, Gerson Lehrman Group, Merck KGaA, Vertex, AstraZeneca, Tango Therapeutics, 3rd Rock, Ono Pharma, Artios, Abingworth, Tesselate, Dark Blue Therapeutics, Pontifax, Astex, Neophore, Glaxo Smith Kline, Dawn Bioventures, Blacksmith Medicines, ForEx, Ariceum. Has stock in: Tango, Ovibio, Hysplex, Tesselate, Ariceum. C.J.L. is also a named inventor on patents describing the use of DNA repair inhibitors and stands to gain from their development and use as part of the ICR “Rewards to Inventors” scheme, and also reports benefits from this scheme associated with patents for PARP inhibitors paid into C.J.L.’s personal account and research accounts at the Institute of Cancer Research. Other authors declare no competing interests.

(*Supplementary information, including Supplementary Figures and figure legends, Supplementary Movies legends and Supplementary Tables legends, can be found in a separate file*)

