## Supplementary Figures and Materials for "Nucleophagy removes cytotoxic trapped PARP1"

*Hoslett et al.*

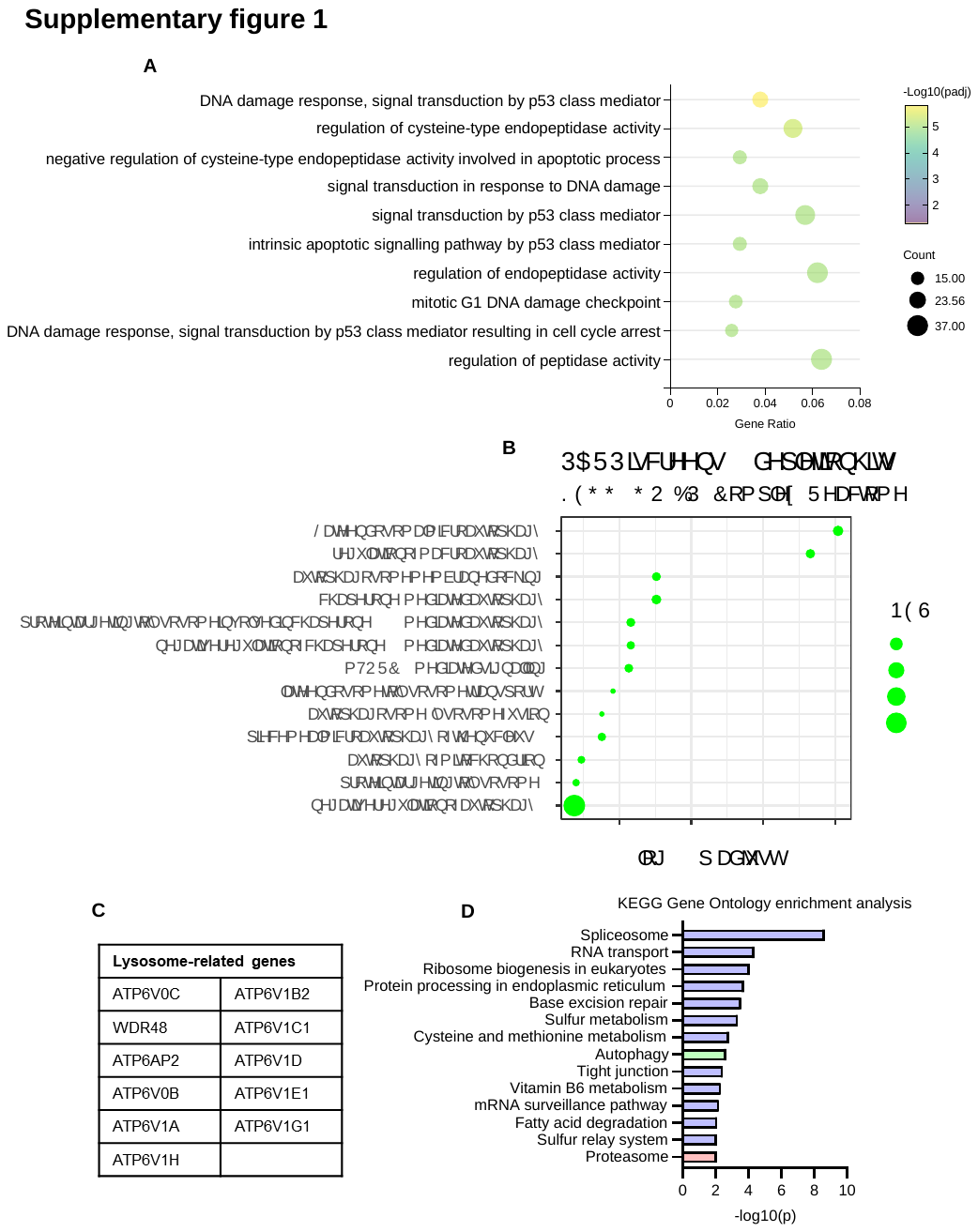

**Supplementary figure 1: Functional pathway analysis in PARPi response. (A)** Gene set enrichment analysis from RNA-seq of genes differentially expressed after treatment with talazoparib (100 nM, 24 hrs). Significance (adjusted p value) is indicated by colour, and the number of genes differentially expressed (count) by bubble size. **(B)** Functional enrichment analysis combined from two whole genome PARPi CRISPR screens, shown in Fig. 1B^31, 76^. **(C)** List of lysosome-related genes shown by blue dots in Fig. 1B. **(D)** Gene set enrichment analysis from mass spectrometry performed by Krastev et al (2022)^39^ of proximity labelling around PARP1. Terms are ranked from top to bottom by significance, with autophagy appearing 8th and proteasome appearing 14th.

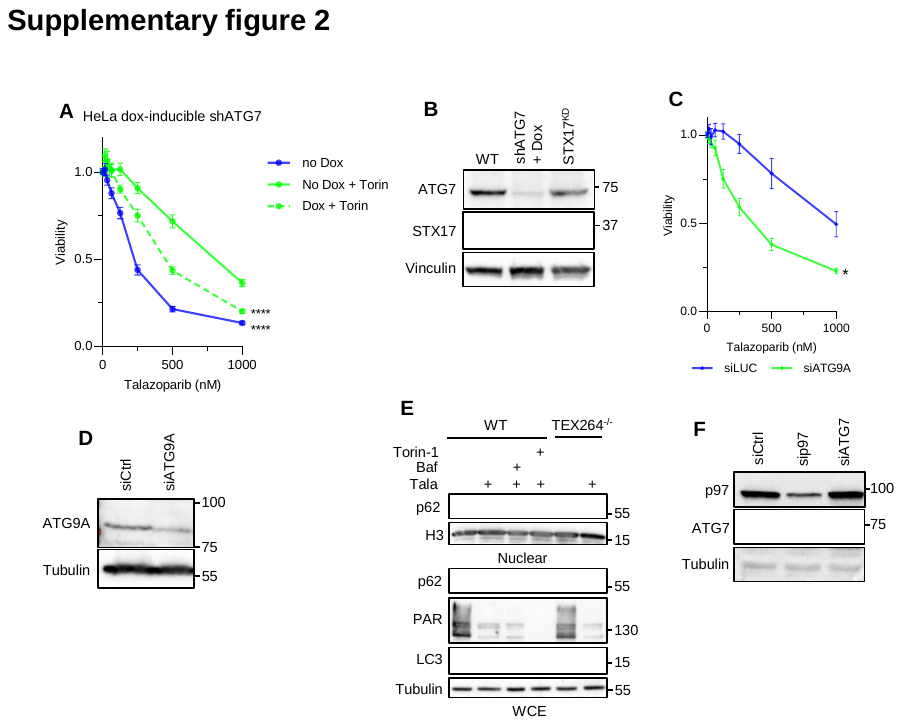

**Supplementary figure 2: (A)** Cell viability measured by resazurin assay in dox-inducible shATG7 HeLa cells treated with talazoparib with or without torin-1 (150 nM) for 24 hrs, followed by 48 hrs recovery. **(B)** Immunoblotting validating depletion of indicated proteins for fig 1E and F. Cell lines are HeLa WT (left lane), HeLa doxycycline-inducible shATG7 (middle lane) and HeLa STX17^KD^ (right lane). **(C)** Cell viability measured by resazurin assay in HeLa cells with depletion of ATG9A using siRNA, treated with talazoparib for 24 hrs, followed by 48 hrs recovery. **(D)** Immunoblot validating depletion of ATG9A in Fig. C. **(E)** Cellular fractionation of HeLa WT and TEX264^-/-^ cells treated with the indicated drugs for 3 hrs. **(F)** Immunoblotting validating depletion of p97 and ATG7 for experiments in Fig. 1 G and H.

**Supplementary figure 3:** **(A)** LysoIP in CAL51 cells stably expressing either PARP1^WT^ or DNA-binding mutant PARP1del.p.119K120S (PARP1^KS^). **(B)** Quantification of (A) from 7 biological repeats. Lysosomal PARP1 signal intensity is normalised by dividing by the input PARP1 signal and lysoIP HA signal. **(C)** LysoIP in HeLa cells treated with the indicated drugs, including niraparib (500 nM) for 3 hrs. **(D)** Quantification of PARP1 levels in (C), normalised to bafilomycin A1-treated. **(E)** Immunoblot showing the expression of mCherry-PARP1-GFP. **(F)** LysoIP in HeLa cells with overexpression of either RNF4 WT (RNF4^WT^) or RNF4 dominant negative M136A+R177A mutant (RNF4^DN^). **(G)** Quantification of PARP1 levels in (F), normalised to the treated control. **(H)** LysoIP in HeLa cells depleted of UFD1 using two different siRNA sequences. **(I)** Quantification of PARP1 levels in (H), normalised to the treated control. All graphs show quantification from at least 3 biological repeats with statistical analysis by unpaired t-test for (B) and (D), and one-way ANOVA for (G) and (I).

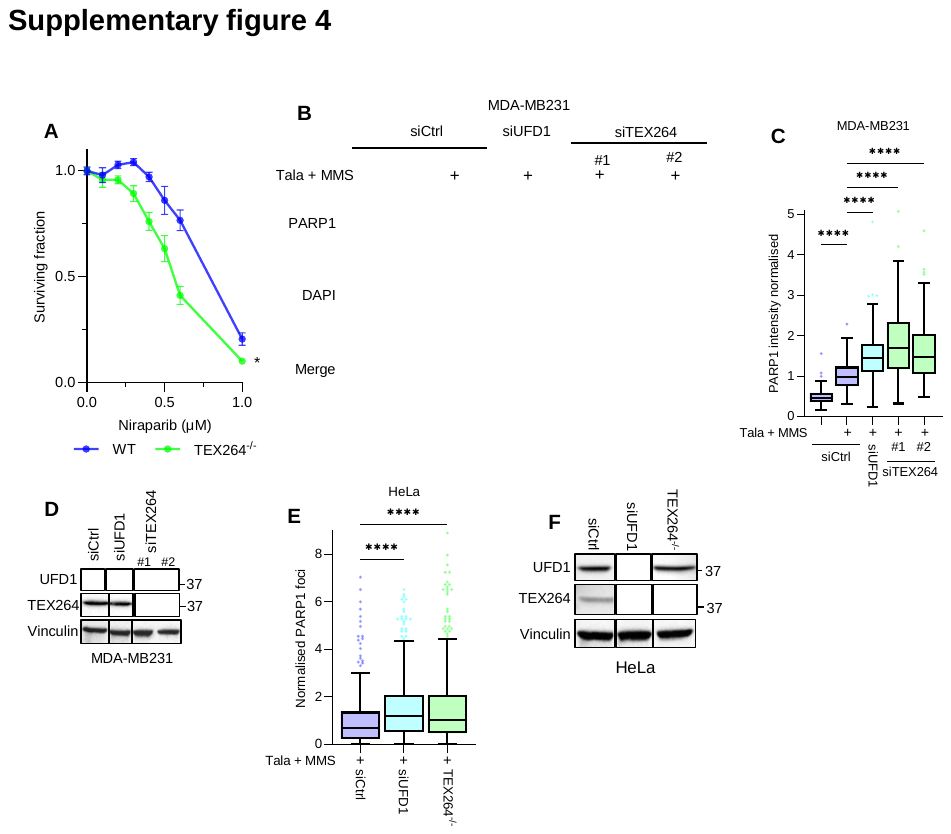

**Supplementary figure 4: (A)** Colony formation assay in HeLa WT and TEX264^-/-^ cells treated with niraparib for 24 hrs. Statistical analysis by two-way ANOVA. **(B)** Images of trapped PARP1 as detected by pre-extraction immunofluorescence in MDA-MB231 cells depleted for the indicated proteins and treated with talazoparib and MMS. **(C)** Quantification of trapped PARP1 levels in (B) from >150 cells in 2 biological repeats, with statistical analysis by one-way ANOVA. **(D)** Validation of depletion for (B and C) by immunoblotting. **(E)** Quantification of trapped PARP1 foci as detected by pre-extraction immunofluorescence in HeLa cells depleted of UFD1 by siRNA or with TEX264^-/-^. Tukey box plot shows >275 cells from 3 biological repeats with statistical analysis by one-way ANOVA. **(F)** Immunoblotting validating knockdown or knockout of proteins in (E).

**Supplementary figure 5: (A)** *In vitro* pulldown of PARP1-GFP combined with TEX264, showing interaction between the two proteins. **(B)** Hydrogen-Deuterium eXchange (HDX-MS) data showing changes in deuterium exchange along the length of TEX264 when combined with PARP1 for 30 s, 10 mins or 60 mins. The green boxes below highlight potential interacting regions. **(C)** *In vitro* pulldown of PARP1-GFP combined with TEX264^WT^ (variant 1) or variants including either the C-terminal (2) or N-terminal (3) regions. Variants 1 and 2 interact strongly, whilst variant 3 does not. **(D)** Co-immunoprecipitation of GFP from the chromatin fraction of HeLa cells stably expressing PARP1-GFP after treatment with talazoparib and MMS for 3 hrs. Cells expressed either TEX264^WT^ or a variant lacking the α-helix region indicated in the schematic (above). The schematic shows the alignment of this variant to sites of interest highlighted by HDX-MS. **(E)** Control conditions for proximity ligation assay between GFP and V5 in cells stably expressing PARP1-GFP and either TEX264-V5 or empty vector (EV)-V5, shown in Fig. 5F. Either or both anti-GFP or anti-V5 antibodies are used as indicated. Scale bar is 10 µm. **(F)** Immunoblot validating the stable expression of TEX264 variants in TEX264^-/-^ CAL51 cells used in Fig. 5J.

**Supplementary figure 6: (A)** PLA between GFP and V5 in HeLa cells stably expressing Lamin-GFP and transiently expressing TEX264-V5. The PLA reaction was performed with anti-GFP and anti-V5 primary antibodies, where indicated, showing the specificity of the PLA signal for TEX264 and Lamin in close proximity. **(B)** LysoIP in HeLa cells treated with the indicated drugs for 3 hrs, including nuclear pore inhibitor leptomycin B (10 nM). **(C)** Quantification of PARP1 levels in (B), normalised to treated control. **(D)** LysoIP in HeLa cells treated with the indicated drugs for 3 hrs, including ATM inhibitor VE-822 (1 µM). **(E)** Quantification of PARP1 levels in (D), normalised to the treated control. Graphs show 3 biological repeats with statistical analysis by students’ unpaired t-test.

**Supplementary figure 7: (A)** Volcano plot showing differential gene expression in HeLa (left) or CAL51 (right) by RNA-seq. Comparison is between TEX264^-/-^ and WT cells after treatment with talazoparib (100 nM, 24 hrs). Genes relating to DNA damage response are coloured as shown. TEX264 is labelled. The grey line shows the adjusted p-value of 0.05. **(B)** Immunoblotting of replication stress markers in HeLa WT and TEX264^-/-^ cells treated with talazoparib for 24 hrs. **(C)** RPA, 53BP1 and γH2AX foci in HeLa WT and TEX264^-/-^ cells after 24 hrs talazoparib treatment. **(D)** Quantification of foci in (C) from >700 cells across 3 biological repeats, shown as a Tukey box plot with statistical analysis by one-way ANOVA. **(E)** γH2AX and RPA foci in CAL51 WT and TEX264^-/-^ cells treated with either talazoparib or veliparib (10µM) for 24 hrs. **(F)** Quantification of foci in (E) in >900 cells from 3 biological repeats as Tukey box plot with statistical analysis by one-way ANOVA. Scale bar on all images is 10 µm.

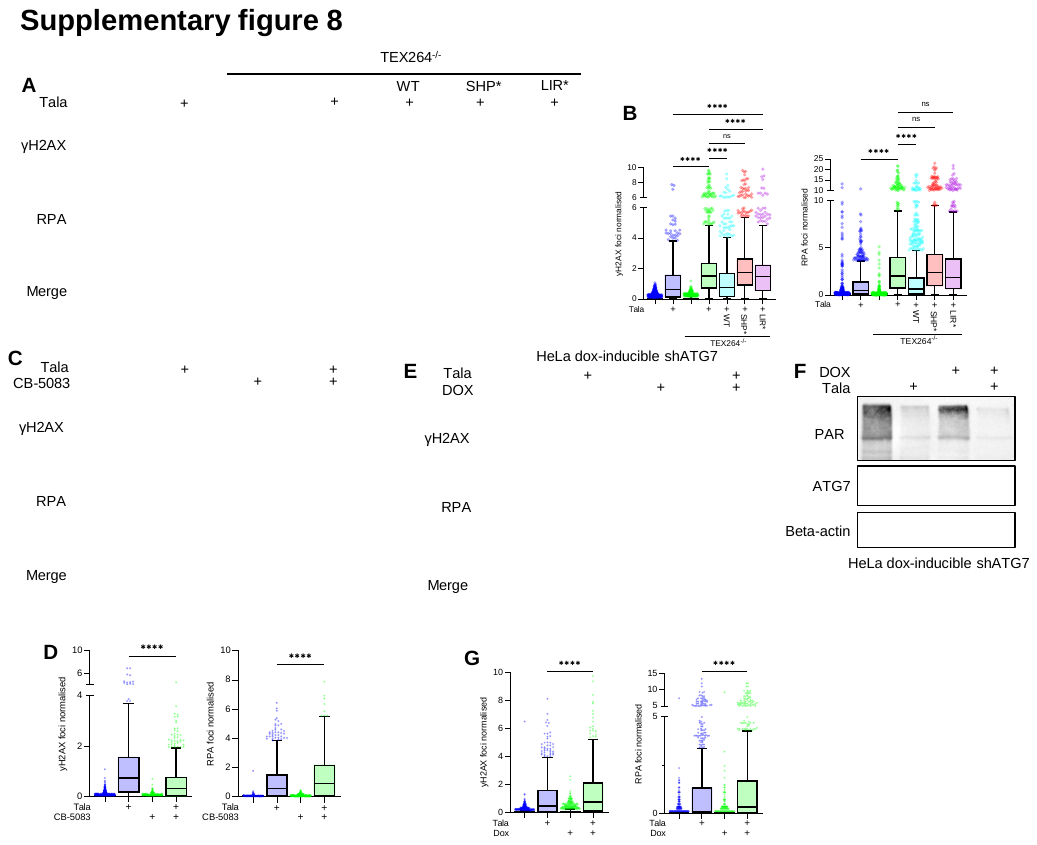

**Supplementary figure 8: (A)** γH2AX and RPA foci in HeLa WT, TEX264^-/-^ or TEX264^-/-^ cells stably expressing indicated TEX264 variants, treated with talazoparib for 24 hrs. (B) Quantification of foci in (A) from >600 cells over 4 biological repeats. **(C)** γH2AX and RPA foci in HeLa WT cells treated with talazoparib for 24 hrs with or without CB-5083 (10 µM) for the last 4 hrs. **(D)** Quantification of foci in (C) from >900 cells over 3 biological repeats as Tukey. **(E)** γH2AX and RPA foci in HeLa doxycycline-inducible shATG7 cells treated with talazoparib for 24 hrs with or without doxycycline. **(F)** Immunoblot validating the reduction in PAR and ATG7 levels when doxycycline-inducible shATG7 cells are treated with talazoparib and doxycycline, respectively. **(G)** Quantification of foci in (E) from >900 cells over 3 biological repeats. All quantifications are shown as Tukey box plots with statistical analysis by one-way ANOVA.

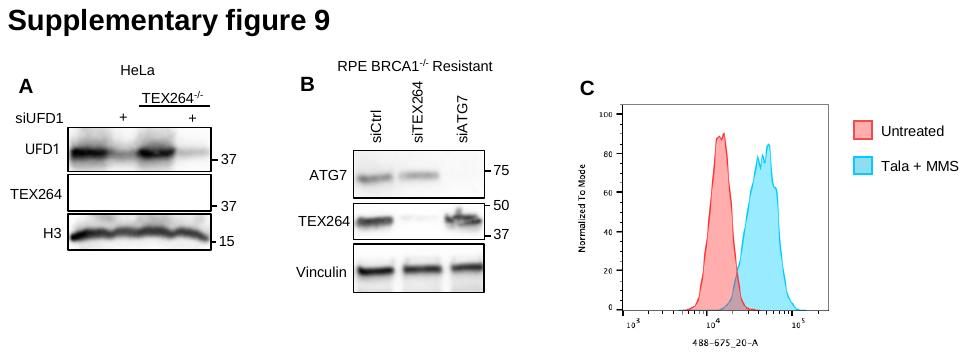

**Supplementary figure 9:** **(A)** Immunoblot validating depletion of UFD1 and TEX264 with siRNA for experiments in Fig. 6A and 6B. **(B)** Immunoblot validating depletion of ATG7 and TEX264 with siRNA for experiments in Fig. 6F and 6G. **(C)** A representative FACS plot from experiments in Fig. 6C showing the increase in proteostat signal in cells treated with talazoparib and MMS for 2 hours, followed by talazoparib for 18 hrs.

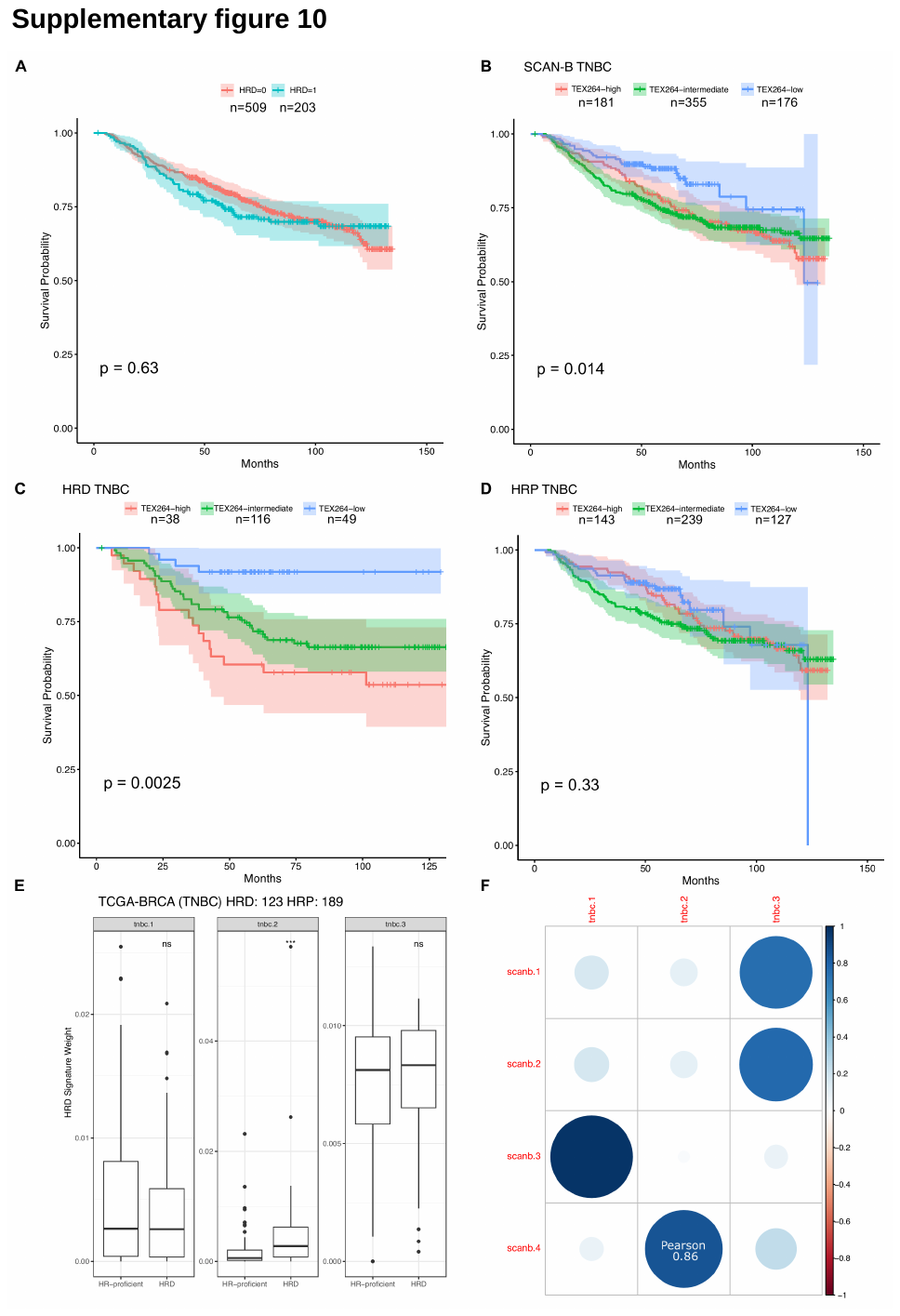

**Supplementary figure 10: (A)** Kaplan-Meier plot of HRD (HRD=1) compared with HRP TNBC from the SCAN-B cohort. **(B)** Kaplan-Meier plot of SCAN-B TNBC by *TEX264* expression based on the highest, lowest quartiles and intermediate expression. **(C)** Kaplan-Meier plot of HRD TNBC based on *TEX264* expression. **(D)** Kaplan-Meier plot of HRD TNBC based on *TEX264* expression. **(E)** Boxplot of HRD signature weights in the TCGA-BRCA TNBC with tnbc.2 correlated with HRD status (Wilcoxon p = 0.00053). **(F)** Correlation matrix of extracted HRD gene expression signatures of the SCAN-B cohort and TCGA-BRCA with scanb.4 matching the HRD signature (tnbc.2) identified in the TCGA-BRCA cohort.

**Movie 1:** PARP1 exists in the nucleus after talazoparib and MMS treatment and is processed by the lysosomes. mCherry-GFP tagged PARP1 monitored by live imaging over 50 minutes by confocal microscopy. mCherry (red), GFP (green) and lysoView 680 (cyan). Timestamps show time since the beginning of the assay. Talazoparib and MMS treatment was added at 2:30. A red puncta can be seen emerging from the nucleus within 5 mins of treatment and is rapidly engulfed by lysosomes, with it mostly degraded by 30 minutes. Scale bar is 5 μm.

**Movie 2:** Rendering of the images shown in movie 1. mCherry (magenta), GFP (green) and lysoView 680 (cyan). Timestamps show time since the beginning of the assay. Talazoparib and MMS treatment was added at 2:30.

**Supplementary Table S1:** Gene Ontology terms from gene set enrichment analysis by RNA-seq in CAL51 talazoparib-treated vs untreated cells, shown in Fig. S1A.

**Supplementary Table S2:** RNA-seq in CAL51 Tala vs untreated cells as shown in Fig. 1A. Autophagy-related genes (from Bordi *et al*, 2021^68^) highlighted in grey. Genes related to positive regulation of the proteasome (GO:1901800) are highlighted in red.

**Supplementary Table S3:** HDX summary table for data shown in fig 5D. The table includes key results for the HDX-MS experiment in both the apo- and holo- state of TEX264 in the presence or absence of PARP1, including the number of peptides identified, sequence coverage, peptide redundancy, and experimental reproducibility ^136^.

**Supplementary Table S4:** HDX uptake summary for the peptides detected in HDX-MS shown in fig 5D.

**Supplementary Table S5:** RNA-seq in HeLa TEX264^-/-^ vs WT cells for Volcano plot in fig S7A. Genes of interest highlighted in accordance with colours in volcano plot.

**Supplementary Table S6:** RNA-seq in CAL51 TEX264^-/-^ vs WT cells for Volcano plot in fig S7A. Genes of interest highlighted in accordance with colours in volcano plot.
